# Stability of the Retinoid X Receptor-alpha Homodimer in the Presence and Absence of Rexinoid and Coactivator Peptide

**DOI:** 10.1101/2020.10.21.333849

**Authors:** Zhengrong Yang, Donald D. Muccio, Nathalia Melo, Venkatram R. Atigadda, Matthew B. Renfrow

**Author notes:** **Corresponding Authors**: Zhengrong Yang - Departments of Biochemistry & Molecular Genetics, University of Alabama at Birmingham, Birmingham, AL, USA 35294; Phone:;, Matthew B. Renfrow - Departments of Biochemistry & Molecular Genetics, University of Alabama at Birmingham, Birmingham, AL, USA 35294; Phone.

## Abstract

Differential scanning calorimetry and differential scanning fluorimetry were used to measure the thermal stability of human retinoid X receptor-alpha ligand binding domain (RXRα LBD) homodimer in the absence or presence of rexinoid and coactivator peptide, GRIP-1. The *apo*-RXRα LBD homodimer displayed a single thermal unfolding transition with a *T*_m_ of 58.7 °C and an unfolding enthalpy (Δ*H*) of 673 kJ/mol (12.5 J/g), much lower than average value (35 J/g) of small globular proteins. Using a heat capacity change (Δ*C_p_*) of 15 kJ/(mol·K) determined by measurements at different pH values, the free energy of unfolding (Δ*G*) of the native state was 33 kJ/mol at 37 °C. Rexinoid binding to the *apo*-homodimer increased *T*_m_ by 5 to 9 °C, and increased the Δ*G* of the native homodimer by 12 to 20 kJ/mol at 37 °C, consistent with the nanomolar dissociation constant (*K*_d_) of the rexinoids. The increase in Δ*G* was the result of a more favorable entropic change due to interactions between the rexinoid and hydrophobic residues in the binding pocket, with the larger increases caused by rexinoids containing larger hydrophobic end groups. GRIP-1 binding to *holo*-homodimers containing rexinoid resulted in additional increases in Δ*G* of 14 kJ/mol, a value same for all three rexinoids. Binding of rexinoid and GRIP-1 resulted in a combined 50% increase in unfolding enthalpy, consistent with reduced structural fluidity and more compact folding observed in other published structural studies. Thermodynamic analysis thus provided a quantitative evaluation of the interactions between RXR and its agonist and coactivator.

## INTRODUCTION

Human Retinoid X Receptors (RXRα, RXRβ, and RXRγ) play a key role in many signaling processes which control cellular proliferation, differentiation, and growth in epithelial tissue as well as maintain proper lipid and glucose homeostasis (1). The three RXR isoforms each display different functions in modulating gene transcription of target genes (2). The proteins are approximately 51 kDa and contain five structural domains. The DNA binding domain (DBD) binds to specific DNA sequences at the start of target genes (3) using two Zn-finger motifs. The ligand binding domain (LBD) at the c-terminal end of RXR is responsible for ligand recognition as well as the associated conformational changes that promote recruitment of coactivator proteins (with replacement of corepressor proteins) important for stabilizing RNA polymerase II to start transcription. The LBD of RXR is connected to the DBD by a short unstructured linker region largely separating the two functions of RXR: specific recognition of target genes and ligand-induced activation of the target gene. Both DBD and LBD contribute to the dimerization interface of the receptor.

The X-ray crystal structures of RXR LBD homodimers alone (*apo*-RXR LBD) or bound to a variety of ligand agonists (*holo*-RXR LBD) revealed the significant conformational changes that allow for agonist induced transcriptional activation (4–7). In *apo*-RXRα LBD, helix 12 extends away from the core of the domain (4), whereas in *holo*-RXRα LBD bound with RXR agonists, helix 12 reorients and folds back toward the ligand binding pocket (LBP) (6). Although these structural changes occur in crystal structures, nuclear magnetic resonance, hydrogen-deuterium exchange mass spectrometry (HDX-MS) and fluorescence polarization studies strongly support that helix 12 is dynamic in both *apo*- and *holo*-homodimers (8–12). The LBD reorganizes around agonists and substantially stabilizes the helical structure at the carboxyl terminus of the LBD, especially helix 11, but not helix 12. Recent X-ray and HDX-MS studies (13–17) of nuclear receptors heterodimers containing the entire RXR bound to double stranded DNA with a target recognition site support the validity of that structural and dynamical conclusions drawn from studies on *holo*-RXRα LBD homodimers.

The recruitment of coactivator proteins to the surface of *holo*-RXR bound to agonist is a key aspect of agonist-induced gene transcription. The steroid receptor coactivator (SRC) family is the most common coactivator proteins of RXR (18, 19). Members of this family have one or more LXXLL sequences that form a putative amphipathic helix which binds to a hydrophobic crevice on the LBD of RXR and nuclear receptors in general. Using coactivator peptides containing the leucine rich motif, numerous high-resolution structures of *holo*-RXR LBD bound to agonist and coactivator peptides have been solved (6, 11–12, 20–24). These structural studies reveal the molecular interactions that occur between the LXXLL motif and helices 3, 4, and 12, which stabilize the active conformation only observed in the presence of agonists. The tertiary structural changes induced by GRIP-1 binding also extend to helix 11 residues that interact directly with the agonists.

Our group has designed and studied many rexinoids for the use in cancer treatment and prevention (23–28). UAB30 (Fig 1B) is a novel rexinoid developed by our group currently in phase II clinical trials. Compared to Targretin, the only clinically used rexinoid approved by the FDA (29), UAB30 exhibits high efficacy in cancer prevention without inducing hyperlipidemia which is the dose-limiting toxicity of Targretin (26, 30–33). UAB110 and UAB111 (Fig. 1B) are two rexinoids belonging to another structural class of UAB rexinoids other than UAB30. Each of these three UAB rexinoids bind the RXRα LBD with nanomolar affinities. UAB110 and UAB111 exhibit potencies approximately 40-fold higher than UAB30 in RXR transcriptional activation (24). However, UAB111 induces hyperlipidemia at its effective dose, while UAB110 does not increase triglyceride levels.

**Figure 1.**
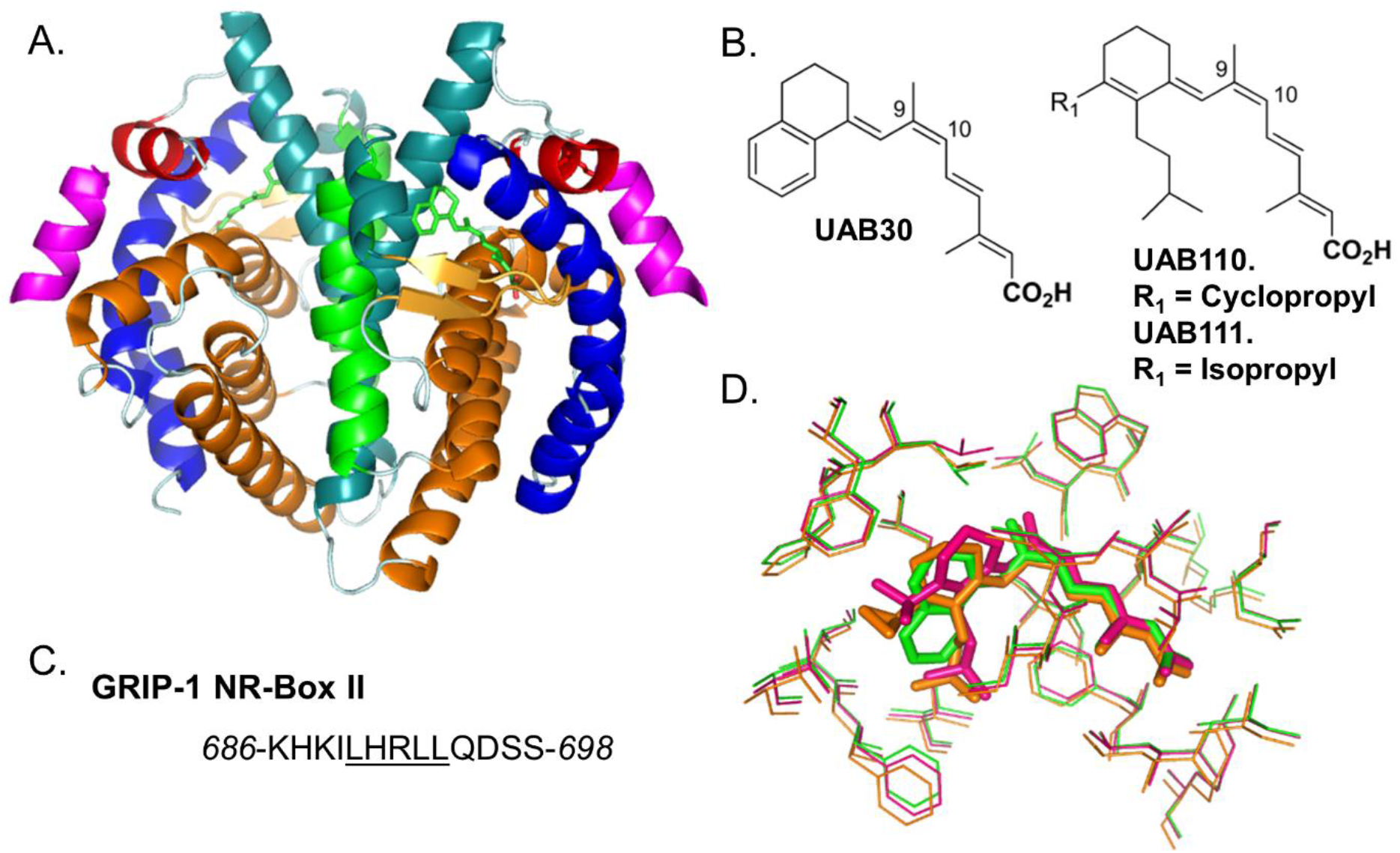
Structures of RXRα LBD, rexinoids and coactivator peptide. A) Crystal structure of RXRα LBD homodimer in complex with UAB30 and GRIP-1 (PDB ID: 4K4J), rendered in Pymol. Secondary elements are colored according to their positions in the three-layer helical sandwich in each monomer. Helices 7 are colored green. Helices 12, which change conformation upon GRIP-1 binding, are colored red. GRIP-1 peptides are colored pink. UAB30 rexinoids are displayed as green sticks. B) Chemical structures of UAB30, UAB110, and UAB111, rendered by ChemDraw. C) The peptide sequence of GRIP-1 NR-Box II coactivator peptide. D) The conformations of UAB30 (green), UAB110 (orange), and UAB111 (deep pink) in the ligand binding pocket of RXRα LBD. The rexinoids are displayed as sticks. The amino acid residues lining the pocket are displayed as thin lines.

To design more potent and nontoxic rexinoids with greater clinical potential and selectivity, we need to fully understand the interactions between the rexinoids and RXRα LBD, and their effects on the structure and dynamics of the RXR homodimers and heterodimers with other nuclear receptors. X-ray crystal structures of RXRα LBD in complex with each UAB rexinoid and a coactivator peptide, GRIP-1 (e.g., RXRα LBD:UAB30:GRIP-1 complex in Fig. 1) revealed only subtle structural differences occurred in the LBP, but there was essentially no difference in the overall backbone fold of the homodimer (Fig. 1D) (11–12, 23–24, 34). Using isothermal titration calorimetry (ITC), we showed that GRIP-1 binds to *holo*-RXR LBD homodimers in complex with each UAB rexinoid with nearly identical affinity and similar enthalpic and entropic signatures (11–12, 23–24). The binding thermodynamics of rexinoids to RXR LBD in the absence of coactivator peptide haven’t been determined, because of limitations to the applicability of ITC in such systems of high affinity, hydrophobic ligands with small binding enthalpy changes (35, 36). Differential scanning calorimetry (DSC) provides an alternative method to obtain thermodynamic data on ligand binding. From the differences in the unfolding temperature and enthalpy with and without a ligand, the thermodynamic parameters of binding of the ligand are determined (37, 38).

Using DSC, we conducted in this study a complete thermodynamic analysis of the thermal unfolding of human RXRα LBD homodimer in the absence or presence of rexinoid and the coactivator peptide, GRIP-1. The goal was to understand how rexinoids and coactivator peptides modulate the energetics of this protein domain, and to gain knowledge on the thermodynamic parameters of interaction between rexinoids and *apo*-RXRα LBD. Differential scanning fluorimetry (DSF), a complementary thermal unfolding method to DSC, was used to inform on key aspects of the unfolding of the domain and to rapidly scan unfolding conditions. We obtained the full panel of unfolding thermodynamic parameters on *apo*-RXRα LBD, including unfolding temperature (*T*_m_), enthalpy (Δ*H*), entropy (Δ*S*), and heat capacity change (Δ*C_p_*^u^). These parameters are essential for analyzing binding energetics of agonists and coactivators to RXR, but have never been reported before. Moreover, from the DSC data obtained in the presence of UAB rexinoids and /or GRIP-1, we calculated the thermodynamic parameters of UAB rexinoids binding to *apo*-RXRα LBD homodimer and GRIP-1 binding to *holo*-RXRα LBD homodimers containing the rexinoids. The latter corresponded closely to those determined by ITC, which demonstrated the validity of this approach. The current study also laid the foundation for future studies on new classes of rexinoids and interaction with heterodimers formed by RXR and other nuclear receptors.

## MATERIALS AND METHODS

### Protein purification

Overexpression and purification of human RXRα LBD (amino acid residues T223-T462) were performed as described by Xia *et al*. (12). Briefly, frozen pellets of BL21-(DE3) *Escherichia coli* bacteria expressing the His6-tagged RXRα LBD fusion protein were lysed by French Press under a pressure of 1250 psi, and centrifuged at 19,000 rpm for 45 min. The clarified supernatant containing the protein was loaded on a 5 mL HiTrap Ni-chelating column (GE Healthcare, Piscataway, NJ), washed, and eluted with an imidazole gradient from 10 to 300 mM in a Tris buffer (20 mM Tris, 500 mM NaCl, pH 8.0). The eluted protein was dialyzed in 2 L of the SEC buffer (10 mM Tris, 50 mM NaCl, 0.5 mM EDTA, 2 mM DTT, pH 8.0) for >4 h. The His_6_-tag was removed by incubating with human α-thrombin overnight. The complete removal of His_6_-tag was confirmed by MALDI-TOF mass spectrometry. The tagless RXRα LBD was then further purified on a HiLoad Superdex 75 size exclusion column (SEC, GE Healthcare, Piscataway, NJ). The fractions containing RXRα LBD homodimers were pooled. SDS-PAGE and MALDI were used to establish a purity of >95% and a mass of the monomers at 27,283 Da. Native PAGE using these fractions displayed a dimer band. Native PAGE using the fractions that eluted just prior to the dimer displayed predominantly a tetramer band, with a less intense dimer band.

### Fluorescence based binding affinity assay

Dissociation constant (*K*_d_) of the rexinoids binding to *apo*-RXRα LBD was determined in a fluorescence based binding assay as previously described (25). The experiments were carried out in a Cary Eclipse Spectrofluorimeter. Three milliliters of a 100 nM *apo*-RXRα LBD solution in the SEC buffer was transferred into a quartz cuvette and equilibrated at 25 °C. Under stirring, the test rexinoid in methanol was added into the protein in 30 1-μL additions. Decrease in fluorescence emission at 337 nm was recorder after each addition, and percent of fluorescence quenching was calculated. The fluorescence quenching data were plotted in Origin 7 (OriginLab, LLC., USA), and nonlinear least square curve fitting was performed to obtain an apparent *K*_d_ based on the following equation (39):

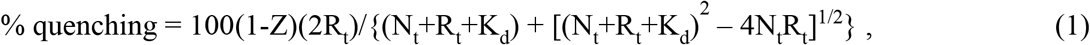

where R_t_ was the total concentration of the test rexinoid, N_t_ was the total concentration of *apo*-RXRα LBD in monomer unit, and Z represented the residual unquenched % fluorescence after the binding site was completely saturated. Nt was determined from the binding curve of UAB111 which had the highest binding affinity. This value was fixed for fitting of UAB30 and UAB110 binding curves.

### Differential scanning calorimetry

Calorimetry experiments were performed using the VP-Capillary DSC System (Malvern Instruments, Westborough, MA) in 0.130 mL cells at a heating rate that varied between of 0.5 and 4 °C/min. An external pressure of 2.0 atm was maintained to prevent possible degassing of the solutions upon heating. Purified *apo*-RXRα LBD homodimer freshly eluted from SEC was dialyzed in 2 L of DSC buffer (10 mM sodium phosphate, 50 mM NaCl, 0.5 mM EDTA, 1 mM TCEP, pH 7.0) for >4 h, and used within 24 h. To collect DSC data at pH values other than 7.0, the purified homodimer was dialyzed in 2 L of borate buffer (10 mM sodium borate, 50 mM NaCl, 0.5 mM EDTA, 1 mM TCEP, pH 8.5 to 9.5) for >4 h, and used within 24 h. To prepared *holo*-RXRα LBD homodimers, rexinoid was added to *apo*-RXRα LBD homodimer from a 100-fold concentrated stock in DMSO. The samples were incubated at 22 °C for 10 min and protected from light. To prepared *holo*-RXRα LBD homodimers bound with coactivator peptide, GRIP-1 was added from a 10 mM stock solution in water. To prepare the buffer control, identical amounts of rexinoid and GRIP-1 were added into the dialysis buffer. Both the protein and the buffer samples were transferred into a 96-well DSC sample loading plate kept in the dark at 5°C and loaded into the DSC cells by the autosampler in the order determined by the sample queue. In some cases, small aliquots of unused *apo*-RXRα LBD homodimer solutions were flash frozen in liquid N2. The aliquots were thawed on ice immediately before sample preparation and used within 24 h. DSC data fitting and analysis were performed using the built-in analysis module in Origin 7 provided by the DSC manufacturer (40). Briefly, after subtraction of the buffer scan, and normalization to molar heat capacity (*C_p_*), a cubic baseline was subtracted from the molar *C_p_* curve to set the pre- and post-transition baselines to zero. The thermal unfolding temperature (*T*_m_) was determined from the maximum of the *C_p_* curve. The calorimetric enthalpy (Δ*H*c) was obtained by integrating the *C_p_* curve. The apparent van’t Hoff enthalpy (Δ*H*v) was obtained by fitting the curve to a build-in model that fits Δ*H*_v_ independently of Δ*H*_c_. The unfolding heat capacity change (Δ*C_p_*^u^) was obtained from the slope of the linear fit of Δ*H*_c_ versus *T*_m_ from data obtained at different pH values.

### Differential scanning fluorimetry

Using the Prometheus NT.48 instrument (NanoTemper Technologies, LLC, South San Francisco, CA) with 48 capillary chambers, each sample was excited at 290 nm and emission was detected simultaneously at 330 and 350 nm. The first derivative of the fluorescence signal at 350 nm relative to 330 nm (F_350_/F_330_) versus temperature produced a DSC-like thermogram. The first derivative of fluorescence intensity at each wavelength exhibited the same thermal unfolding profiles. Because the UV-vis absorption of the rexinoids interfered with the detection of unfolding using F_350_/F_330_, DSF data of fluorescence intensity at 350 nm were used to obtained *T*_m_. Sample preparation for DSF was the same as for DSC. Each capillary required 10 μL to load. Duplicate measurements were performed on each sample. *T*_m_ of each sample was automatically determined by the built-in analysis software and tabulated in an Excel output file.

### Circular dichroism

CD spectra were recorded in a OLIS CD spectrophotometer (OLIS, Athens, GA). Quartz cuvettes with a path-length of 0.02 cm were used. The protein concentration was 12 μM dimer. The CD cell holder was heated using an external water bath to the desired temperatures. CD spectra were recorded from 260 to 190 nm. Buffer baselines were recorded at the same temperatures and subtracted from the protein spectra. Total data collection time was ~5 min for each spectrum. For the completely denatured sample in guanidine-HCl, the sample was prepared by adding guanidine-HCl as a solid into the thermally unfolded protein solution to reach a final concentration of 5 M. The sample was then cooled to 25 °C in the cell holder, and the CD spectra were collected at 25 °C. The final protein concentration was adjusted based on the volume change after the addition of the denaturant. Optical interference from the denaturant prevented scans below 210 nm.

### Isothermal titration calorimetry

ITC experiments were performed on an Auto-iTC200 System (Malvern Instruments, Westborough, MA). For titration of UAB30 with *apo*-RXRα LBD, the ITC sample cell contained 200 μL of 5 μM UAB30, prepared by diluting 500 μM UAB30 in 100% DMSO into the DSC buffer (pH 7.0). The ITC syringe contained 20 μM *apo*-RXRα LBD homodimer in DSC buffer with 1% w/v DMSO. Each titration experiment consisted of 16 injections of 2.5 μL of *apo*-RXRα LBD into UAB30 at 10, 20, or 30 °C. Background mixing heat was determined from injections of *apo*-RXRα LBD into the same buffer without UAB30. For titration of RXRα LBD:rexinoid with GRIP-1, the ITC sample cell contained 12.5 μM RXRα LBD dimer and 35 μM test rexinoid in the DSC buffer with 1% w/v DMSO. The ITC syringe contained 350 μM GRIP-1 in the same buffer. Each titration experiment consisted of 16 injections of 2.5 μL of GRIP-1 into RXRα LBD:rexinoid at 10, 20, or 30 °C. Background mixing heat was determined from injections of GRIP-1 into the same buffer without RXRα LBD. Data analysis was performed using the built-in analysis module in Origin 7 provided by the ITC manufacturer. Monomeric protein concentration was used for analysis to obtain the stoichiometry of binding to each RXRα LBD monomer.

### Calculation of unfolding thermodynamic parameters

The following equations were used to calculate Δ*G*, Δ*H*, and Δ*S* of unfolding as a function of *T* from the experimentally determined *T*_m_, Δ*H*_c_, and Δ*C_p_*^u^:

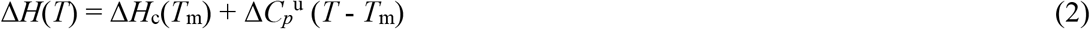

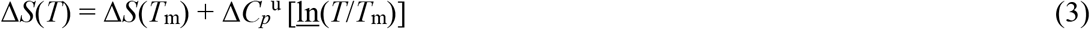

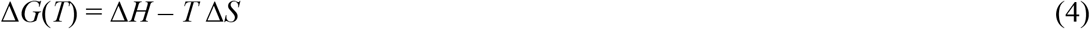

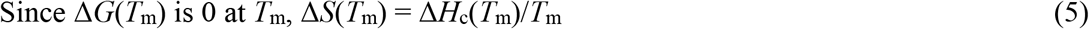

### Calculation of rexinoid binding constant

Binding constant (*K*_a_) of rexinoid to *apo*-RXRα LBD at the unfolding temperature was calculated from the increase in unfolding temperature of apo-RXRα LBD in the presence of rexinoids based on the following equation (41):

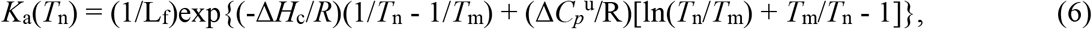

where *T*_n_ was the unfolding temperature of *apo*-RXRα LBD in the presence of rexinoid, *T*_m_ was the unfolding temperature in the absence of rexinoid, *R* was the universal gas constant, Δ*H*_c_ was the calorimetric unfolding enthalpy of *apo*-RXRα LBD in the absence of rexinoid at *T*_m_, Δ*C_p_*^u^ was the unfolding heat capacity change in the absence of rexinoid, and L_f_ was the free rexinoid concentration at *T*_n_, which equaled to total rexinoid concentration minus half of the total protein concentration. Calculation of *K*_a_ was performed on a per monomer basis assuming the two binding sites of the homodimer (one in each monomer) are identical and noncooperative. To minimize unwanted processes due to great excesses of ligand (37), *K*_a_ was calculated based on the shift in *T*_m_ at 1:1 rexinoid/monomer molar ratio.

### Calculation of GRIP-1 binding constant

*K*_a_ of GRIP-1 binding at the unfolding temperature was calculated from the increase in unfolding temperature of holo-RXRα LBD in the presence of GRIP-1 based on Equation (6), where *T*_n_ was the unfolding temperature of *holo*-RXRα LBD in the presence of GRIP-1, *T*_m_ was the unfolding temperature in the absence of GRIP-1, Δ*H*_c_ was the calorimetric unfolding enthalpy of *holo*-RXRα LBD in the absence of GRIP-1 at *T*_m_, Δ*C_p_*^u^ was the unfolding heat capacity change in the absence of GRIP-1, and L_f_ was the free GRIP-1 concentration at *T*_n_, which equaled to total GRIP-1 concentration minus half of the total protein concentration. *K*_a_ of GRIP-1 binding at other temperatures was calculated based on the integrated van’t Hoff equation:

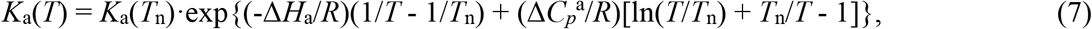

where *R* was the universal gas constant, *T*_n_ was the unfolding temperature of *holo*-RXRα LBD in the presence of GRIP-1, Δ*H*_a_ was the GRIP-1 binding enthalpy at *T*_n_, and Δ*C_p_*^a^ was the binding heat capacity change.

## RESULTS AND DISCUSSION

### I. Thermal Unfolding of *Apo*-RXRα LBD Homodimers

The Δ*G* of *apo*-RXRα LBD unfolding at 30 °C was previously determined by Harder *et al*. (42) using isothermal chemical denaturation. However, the enthalpic and entropic changes of the unfolding process were not obtained, nor was the unfolding heat capacity change which contributes significantly to the temperature dependence of Δ*G*.

#### 1.1 Partial reversibility of transition

To determine the stability of *apo*-RXRα LBD homodimer at 37 °C and other temperatures, thermal unfolding of this homodimer was measured by DSC from 35 to 70 °C using a scan rate (*ν*) of 4 °C/min. A single endotherm was observed centered at the maximum thermal unfolding temperature (*T*_m_) of 58.2 ± 0.1 °C, with a calorimetric unfolding enthalpy (Δ*H*_c_) of 665 ± 4 kJ/mol (Fig. 2, **trace 1**). Upon rescan, no thermal unfolding endotherm was observed, indicating the unfolding of the homodimer was irreversible when heated to 70 °C (Fig. 2, **trace 2**). To examine whether partial reversibility occurred during the unfolding process, DSC endotherms were obtained on homodimers systematically heated for one minute at various temperatures and then rapidly cooled in an ice bath. When the incubation temperature was 57.3 °C, which corresponded to 25% unfolding^1^, an endotherm was observed with the same *T*_m_, but with approximately 80% the intensity of the first scan (Fig. 2, **trace 3**). As the incubation temperature was increased to 58.2 °C (50% unfolding), the Δ*H*_c_ was decreased to 50% (Fig. 2, **trace 4**). The Δ*H*_c_ was further decreased to 30% when the incubation temperature was increased to 60.0 °C (75% unfolding, Fig. 2, **trace 5**). This data supports that the unfolding of the homodimer was partially reversible through the endotherm.

**Figure 2.**
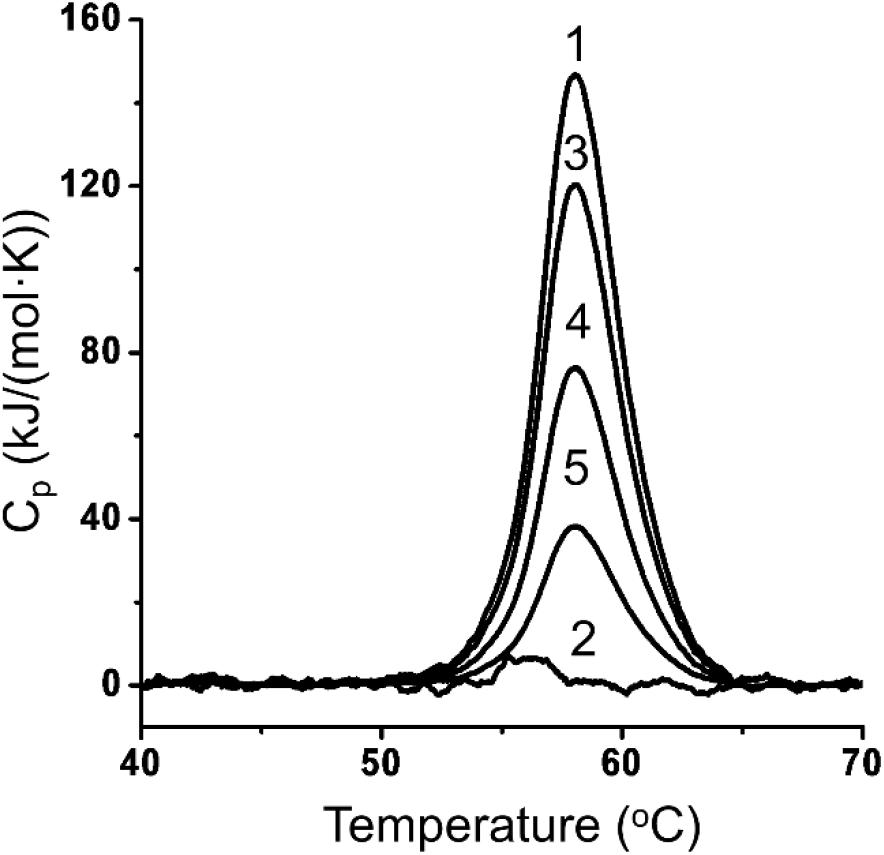
Thermal unfolding of *apo*-RXRα LBD is partially reversible. DSC molar heat capacity profile of 1.5 μM *apo*-RXRα LBD homodimer. (1) First scan; (2) rescan; (3) after heating at 57.3 °C for 1 min; (4) after heating at 58.6 °C for 1 min; (5) after heating at 60.0 °C for 1 min.

The scan rate dependence of the thermal unfolding of *apo*-RXRa LBD was next examined. The scan rate, *v*, of the DSC measurement was decreased from 4.0 to 0.5 °C/min. As displayed in Fig. 3A and summarized in Table 1, *T*_m_ decreased systematically as *v* decreased, reaching 55.6 ± 0.1 °C when *v* was 0.5 °C/min. Δ*H*_c_ remained nearly constant at 670 kJ/mol regardless of *v*. An apparent van’t Hoff enthalpy (Δ*H*_v_) was determined by non-linear least square curve fitting to a two-state model, whereby a natively folded protein, N, unfolds reversibility to U: N ↔ U (43). Δ*H*_v_ systematically increased from 815 ± 13 kJ/mol when *v* was 4 °C/min to 1208 ± 4 kJ/mol when *v* was 0.5 °C/min.

**Table 1.**
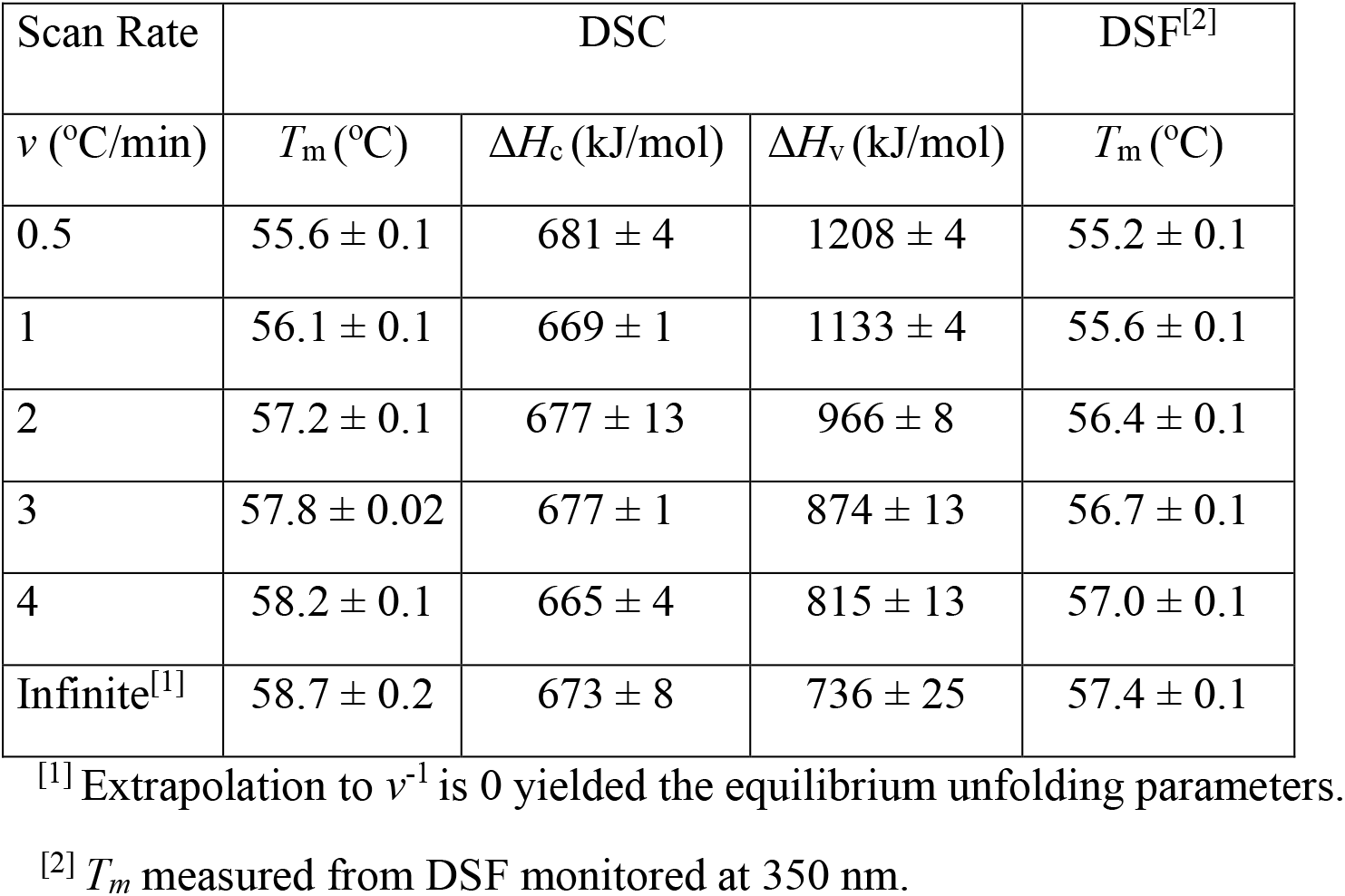
Scan rate dependence of the thermal unfolding parameters of *apo*-RXRα LBD.

| Scan Rate | DSC |  |  | DSF<sup>[2]</sup> |
| --- | --- | --- | --- | --- |
| $\nu$ (°C/min) | $T_m$ (°C) | $\Delta H_c$ (kJ/mol) | $\Delta H_v$ (kJ/mol) | $T_m$ (°C) |
| 0.5 | $55.6 \pm 0.1$ | $681 \pm 4$ | $1208 \pm 4$ | $55.2 \pm 0.1$ |
| 1 | $56.1 \pm 0.1$ | $669 \pm 1$ | $1133 \pm 4$ | $55.6 \pm 0.1$ |
| 2 | $57.2 \pm 0.1$ | $677 \pm 13$ | $966 \pm 8$ | $56.4 \pm 0.1$ |
| 3 | $57.8 \pm 0.02$ | $677 \pm 1$ | $874 \pm 13$ | $56.7 \pm 0.1$ |
| 4 | $58.2 \pm 0.1$ | $665 \pm 4$ | $815 \pm 13$ | $57.0 \pm 0.1$ |
| Infinite<sup>[1]</sup> | $58.7 \pm 0.2$ | $673 \pm 8$ | $736 \pm 25$ | $57.4 \pm 0.1$ |

<sup>[1]</sup> Extrapolation to $\nu^{-1}$ is 0 yielded the equilibrium unfolding parameters.
<sup>[2]</sup> $T_m$ measured from DSF monitored at 350 nm.

**Figure 3.**
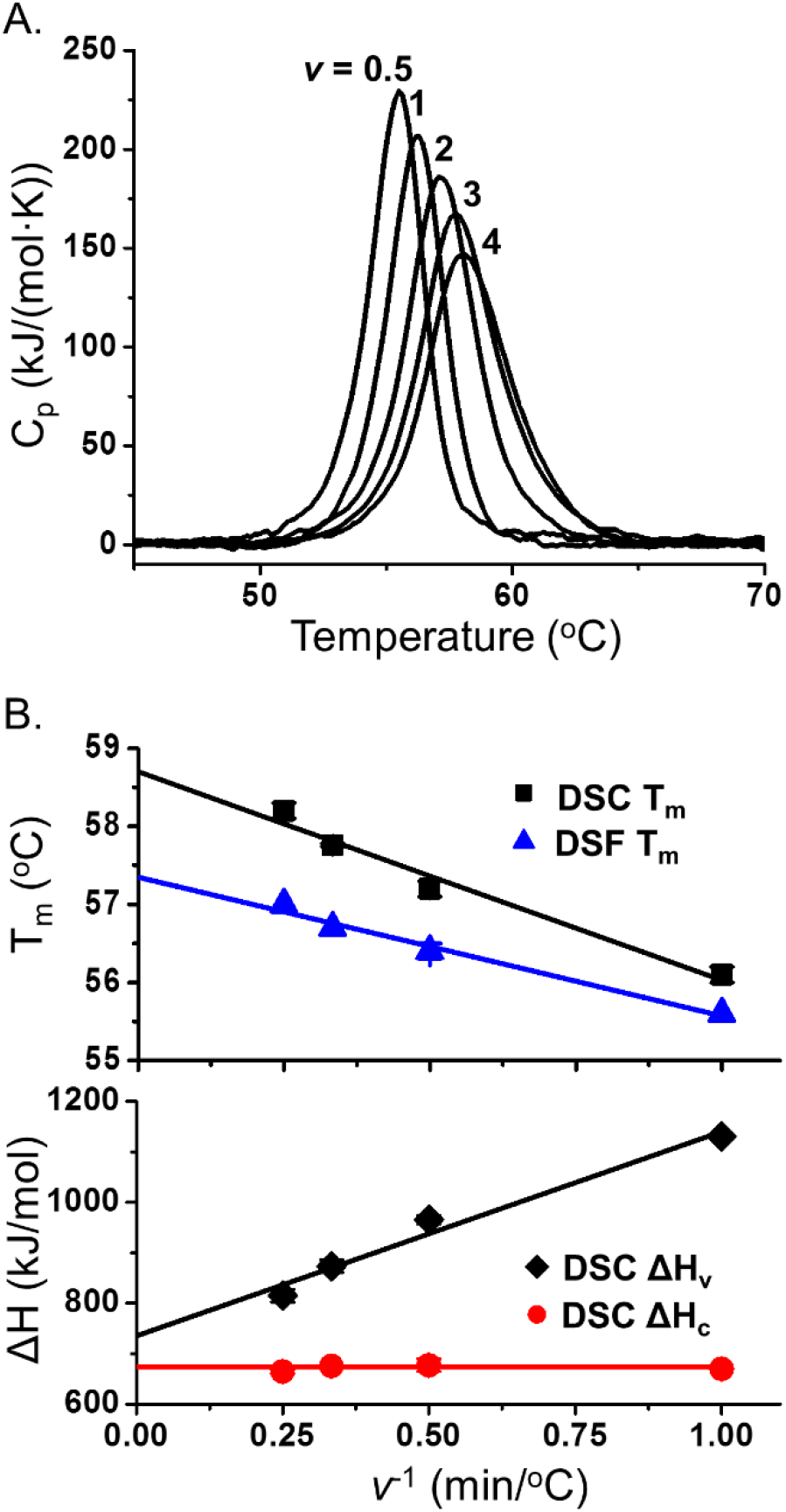
Equilibrium unfolding parameters of *apo*-RXRα LBD obtained by extrapolation to infinite scan rate. A) DSC molar heat capacity profiles for 1.5 μM *apo*-RXRα LBD homodimer at different scan rates, *v* (from left to right): 0.5, 1.0, 2.0, 3.0, and 4.0 °C/min. B) *T*_m_, Δ*H*_c_, and Δ*H*_v_ as a function of *v*^−1^. The *T*_m_ and Δ*H*_v_ values were fitted linearly to *v*^−1^. The equilibrium unfolding parameters were obtained by extrapolation to zero *v*^−1^.

The thermal unfolding of *apo*-RXRα LBD was also monitored by measuring the changes in intrinsic fluorescence at 330 and 350 nm (**Fig. S1**). The changes in tryptophan (Trp) emission intensity at 330 nm or 350 nm reflected the change in Trp environments during unfolding. The differential scanning fluorimetry (DSF) curves of *apo*-RXRα LBD, which contains two Trp residues, displayed only one unfolding transition that was similar to the DSC endotherms. The trends in the *T*_m_ at different scan rates matched those determined by DSC (Table 1), suggesting both DSC and DSF detected the same global unfolding event. The *T*_m_ values determined by DSF were systematically lower than the *T*_m_ measured by DSC by ~1 °C. The lower *T*_m_ measured by DSF relative to those by DSC was observed for other protein unfolding processes (44–46). Trp residues in folded states may sense changes in their local environments at a slightly lower temperature than onset of the global unfolding measured calorimetrically (47).

The lack of reversible unfolding transition in DSC rescans and the dependence of *T*_m_ on *v* indicated that the thermal unfolding of RXRα LBD was not in complete equilibrium throughout the heating process. DSC irreversibility is a common phenomenon, especially for multi-domain or oligomeric proteins (48). Often protein aggregation occurs at higher temperatures, which prevents reversibility (49). However, these irreversible DSC transitions yield equilibrium data if the unfolding follows the Lumry-Eyring model: whereby N unfolds reversibility to U, which converts to a denatured state (D) slowly and irreversibility: N ↔ U → D (50). If the irreversible step occurs slower than the rate of protein unfolding and refolding, then thermodynamic data for the reversible step is obtained by extrapolation of measured data to infinite scan rate (48, 51–53). For *apo*-RXRα LBD, the *T*_m_ and Δ*H*_v_ were plotted versus *v*^−1^ (Fig. 3B); each thermal parameter was linearly dependent of *v*^−1^ for *v* faster than 0.5 C/min. The extrapolated equilibrium unfolding parameters (Table 1) were: *T*_m_^eq^ = 58.7 ± 0.2 °C, Δ*H*_v_^eq^ = 736 ± 25 kJ/mol, and Δ*H*_c_^eq^ = 673 ± 8 kJ/mol (using the average of the apparent Δ*H*_c_ at different *v* values).

#### 1.2 Two-state unfolding without dimer dissociation

The ratio of the van’t Hoff enthalpy to the calorimetric enthalpy, Δ*H*_v_^eq^/Δ*H*_c_^eq^, was essentially one using the extrapolated parameters. A cooperative unit of 1 when both Δ*H*_c_^eq^ and Δ*H*_v_^eq^ were determined based on per mole of dimer indicated that the native dimer was the cooperative unfolding unit. The ratio of 1 for Δ*H*_v_/Δ*H*_c_ was not expected if the dimeric protein dissociated during unfolding: N2 ↔ 2U. For this unfolding equilibrium, the unfolding endotherm is asymmetrical about its midpoint, and the apparent Δ*H*_v_/Δ*H*_c_ is about 0.7 (53) (see **Fig. S2** for a simulated DSC transition based on the N2 ↔ 2U model, in comparison to the N ↔ U model). The asymmetry and low Δ*H*_v_/Δ*H*_c_ was not observed in any of the DSC traces obtained in this study, suggesting that the native dimer unfolded without significant dissociation to monomers (54–56).

To examine this further, native gels were used to determine the aggregation state of the homodimer at three different temperatures: 57 °C which was below *T*_m_, 58.7 °C which was the extrapolated equilibrium *T*_m_, and 65 °C which was above the completion of the endotherm. As displayed in Fig. 4, *apo*-RXRα LBD homodimer migrated near 50 kDa, consistent with a dimer; whereas the tetramer migrated near 100 kDa. When the dimer was heated at 57 °C for 1, 2 and 5 min and rapidly cooled in an ice bath, the band corresponding to the dimer was observed but the intensity decreased as heating time increased. In addition, a species that migrated significantly above that of the unheated dimer emerged and increased in intensity, consistent with an aggregate of much higher molecular weight. When the heating temperature increased to 58.7 °C, less dimer and more aggregated species were observed. When heated to 65 °C for these times, the dimer disappeared and only a highly aggregated species was present in the native gels. A band consistent with monomer was not observed in any lane.

**Figure 4.**
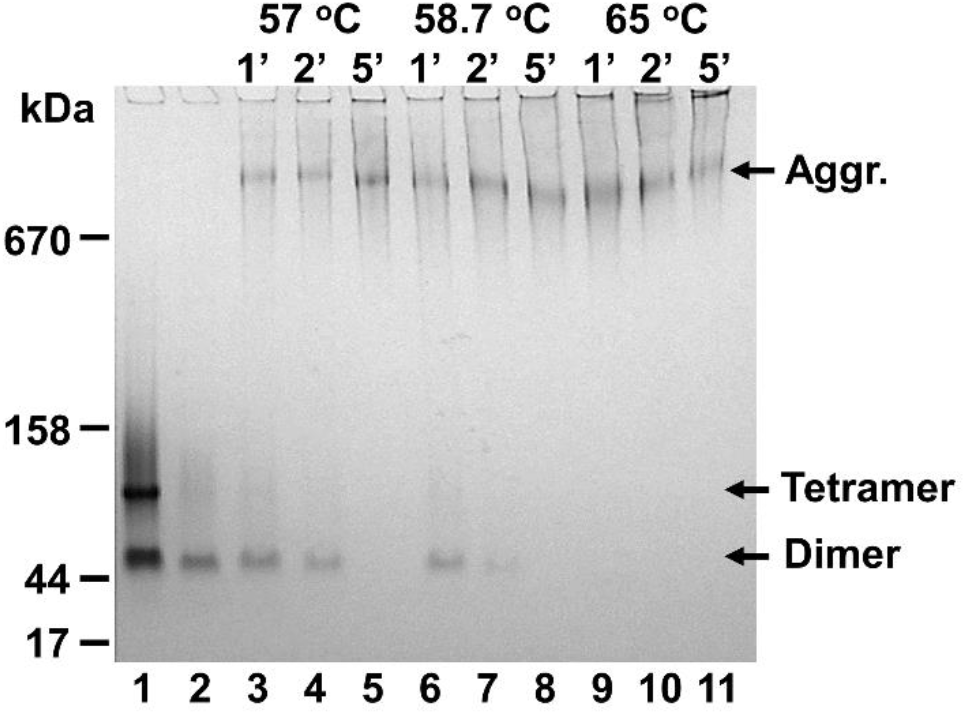
Changes in *apo*-RXRα LBD aggregation state during thermal unfolding. Native PAGE of 5 μM *apo*-RXRα LBD homodimer incubated at different temperatures for 1, 2, and 5 min, and then rapidly cooled in an ice bath. Lane 1: Fraction from SEC containing a mixture of tetramer and dimer that was not heated. Lane 2: Fraction from SEC containing only dimer that was not heated. Lanes 3 to 5: dimer heated at 57°C for 1, 2, and 5 min. Lanes 6 to 8: dimer heated at 58.7°C for 1, 2, and 5 min. Lanes 9 to 11: dimer heated at 65°C for 1, 2, and 5 min.

Circular dichroism (CD) and fluorescence spectroscopy was measured at several temperatures to inform on changes in secondary and tertiary structure during the endotherm. The X-ray crystal structure of *apo*-RXRα LBD contained 12 helices composing about 66% of its secondary structure (4, 7). The 222 and 208 nm negative CD signals (Fig. 5A), which are significant in helical protein structures, were present in CD spectra at temperatures up to 56 °C (12). The magnitude of the 208 and 222 nm negative bands decreased ~20% when heated to temperatures near to *T*_m_ measured by DSC. These signals decreased ~40% when the dimer was heated to 75 °C, slightly higher than the temperature of the end of the DSC endotherm. No further decrease in CD signals was observed when the sample was incubated at 75 °C for 10 more minutes. This indicated that the secondary structure of *apo*-RXRα LBD was not completely unfolded throughout the DSC endotherm. In contrast, the 222 and 208 nm signals were completely lost for the homodimer in the presence of 5 M guanidine-HCl at 25 °C, which completely denatured and dissociated the dimer into monomers (42).

**Figure 5.**
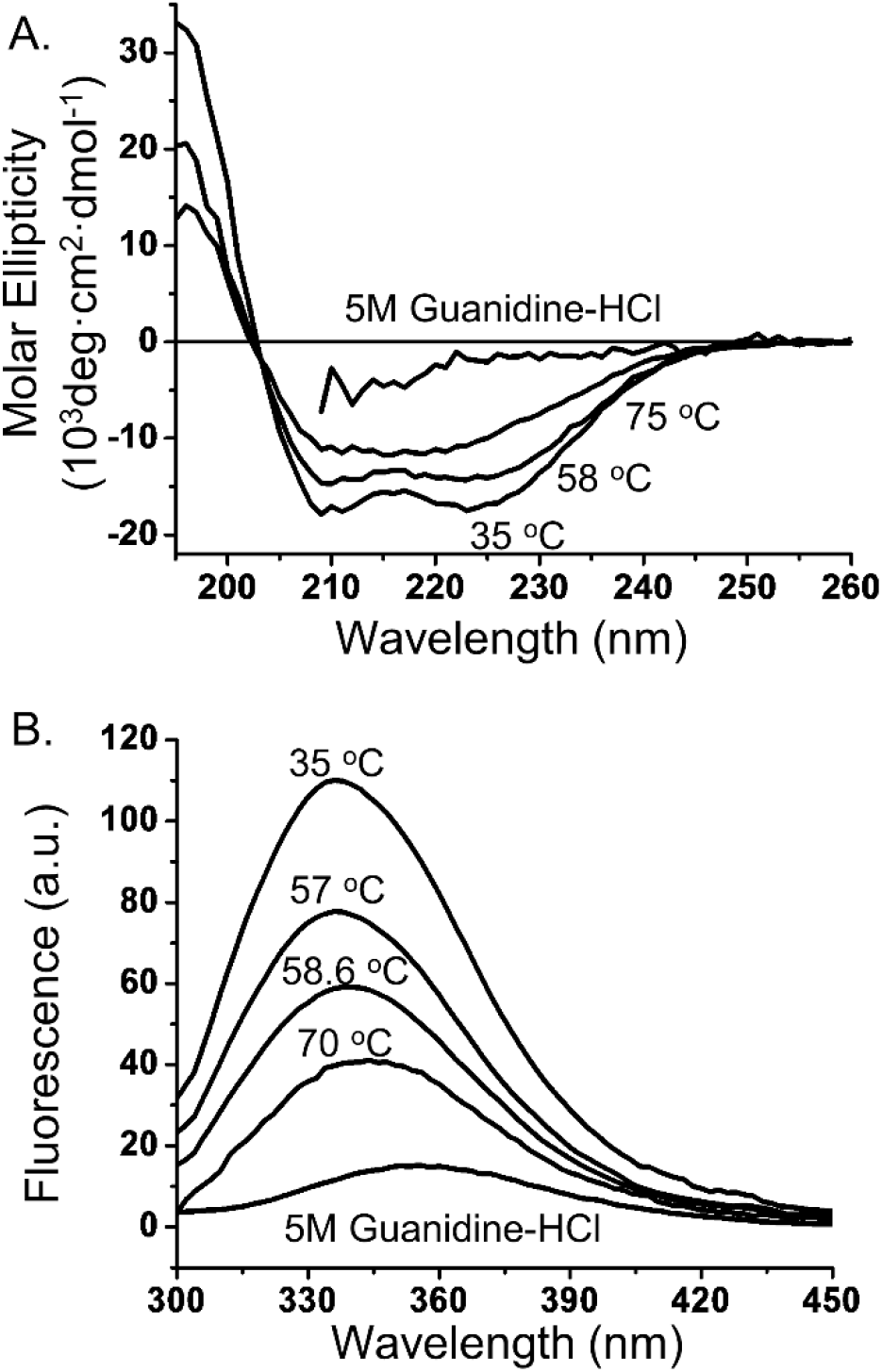
Spectroscopic changes in *apo*-RXRα LBD during thermal unfolding. A) Circular dichroic spectra of 12 μM *apo*-RXRα LBD homodimer at different temperatures and in the presence of 5 M guanidine-HCl at 25 °C. B) Fluorescence spectra of 0.05 μM *apo*-RXRα LBD homodimer at different temperatures and in the presence of 5 M guanidine-HCl at 25 °C. The excitation wavelength was 280 nm.

Each monomer of *apo*-RXRα LBD contains two Trp: W282 in helix 2 and W305 in helix 4, which is near the rexinoid binding site. An intense fluorescence band centered at 335 nm was observed at 35 °C, consistent with the hydrophobic environment of indole group of the two Trp in a folded state (Fig. 5B). Upon heating the protein to higher temperatures, the Trp fluorescence intensity decreased due to thermal quenching (57), and the emission maximum wavelength (λ_max_) gradually red-shifted to 344 nm at 70 °C. A red-shift of λ_max_ to 355 nm was observed for the completely denatured homodimer in 5 M guanidine-HCl. The much less red-shifted signals for the thermally denatured *apo*-RXRα LBD indicated that either one or both of the Trp were still in a partially hydrophobic environment when heated to 70 °C. Taken together, these thermodynamic and spectral data are most consistent with the native homodimer being converted to a partially unfolded and aggregated *apo*-RXRα LBD species above *T*_m_.

#### 1.3 Thermodynamic unfolding parameters at 37 °C

Using a *v* of 4 °C/min for DSC measurements, the time needed to unfold the protein was less than 3 min, which minimized the effects of the irreversible processes during the unfolding transition. This is reflected in the fact that the unfolding parameters obtained at 4 °C/min were very similar to the extrapolated equilibrium parameters (Table 1). Therefore, all subsequent DSC experiments were conducted at 4 °C/min. To modulate unfolding *T*_m_ and enthalpy for determining the unfolding heat capacity change (Δ*C_p_*^u^), the pH of the buffer was changed. Due to rapidity of measurement and lower sample amounts, DSF was first used to survey a range of pH values between 5 to 9 (**Fig. S3**). DSF showed that *apo*-RXRα LBD was most stable at pH 7.0, and it required an increase of 2 pH units to induce significant decreases in *T*_m_. DSC was performed on *apo*-RXRα LBD homodimer at four different pH values: 7.0, 8.6, 9.0, and 9.5 (Fig. 6A). Δ*H*_c_ was fit linearly to *T*_m_ (Fig. 6B). From the slope of the line, Δ*C_p_*^u^ was determined to be 15 ± 1 kJ/(mol·K). Using this Δ*C_p_*^u^ and the extrapolated values of *T*_m_ and Δ*H*_c_ from the DSC measurements (Table 1), Δ*H* of unfolding for the *apo*-RXRα LBD homodimer was calculated to be 347 ± 21 kJ/mol at 37 °C. Δ*S* of unfolding was 1.01 ± 0.06 kJ/(mol·K), −*T*Δ*S* was −314 ± 17 kJ/mol, and Δ*G* of unfolding was 33 ± 3 kJ/mol at 37 °C (Table 2). A Δ*G* of 32 ± 3 kJ/mol was calculated using parameters obtained at a *v* of 4 °C/min, which was within experimental error of the Δ*G* calculated from the extrapolated parameters. *Apo*-RXRα LBD homodimer was most stable at 20 °C, with a Δ*G* of 43 ± 3 kJ/mol.

**Table 2.** Thermodynamic parameters of unfolding of *apo*-RXRα LBD, *holo*-RXRα LBD with and without GRIP-1.

| | $T_m$ | $\Delta H_c$ | $\Delta H_v$ | $\Delta C_p^u$ | $\Delta H$<br>(37 °C) | $-T\Delta S$<br>(37 °C) | $\Delta G$<br>(37 °C) |
| --- | --- | --- | --- | --- | --- | --- | --- |
|  | (°C) | (kJ/mol) | (kJ/mol) | (kJ/<br>(mol·K)) | (kJ/mol) | (kJ/mol) | (kJ/mol) |
| *Apo*-RXRα LBD <sup>[1]</sup> | 58.7 ± 0.2 | 673 ± 8 | 736 ± 25 | 15 ± 1 | 347 ± 21 | -314 ± 17 | 33 ± 3 |
| DMSO control <sup>[2]</sup> | 57.9 ± 0.4 | 690 ± 8 | 803 ± 4 | 15 ± 1 | 376 ± 17 | -342 ± 17 | 33 ± 2 |
| + GRIP-1 | 59.3 ± 0.2 | 690 ± 29 | 811 ± 13 | 14 ± 2 | 389 ± 52 | -353 ± 48 | 36 ± 7 |
| + UAB30 | 63.6 ± 0.1 | 803 ± 4 | 836 ± 13 | 17 ± 1 | 371 ± 26 | -326 ± 23 | 45 ± 4 |
| + UAB110 | 66.6 ± 0.5 | 849 ± 13 | 811 ± 17 | 16 ± 2 | 362 ± 37 | -310 ± 32 | 52 ± 8 |
| + UAB111 | 66.9 ± 0.3 | 840 ± 21 | 823 ± 21 | 17 ± 1 | 362 ± 24 | -309 ± 20 | 53 ± 5 |
| + UAB30/GRIP-1 | 67.5 ± 0.3 | 932 ± 33 | 924 ± 25 | 17 ± 1 | 374 ± 35 | -317 ± 30 | 58 ± 8 |
| + UAB110/GRIP-1 | 69.9 ± 0.5 | 995 ± 17 | 924 ± 8 | 18 ± 1 | 393 ± 35 | -327 ± 30 | 66 ± 8 |
| + UAB111/GRIP-1 | 70.4 ± 0.2 | 991 ± 29 | 928 ± 8 | 20 ± 2 | 380 ± 35 | -314 ± 30 | 66 ± 9 |

<sup>[1]</sup> Equilibrium unfolding parameters obtained by extrapolating to $v^{-1}$ is zero, in the absence of DMSO.
<sup>[2]</sup> DMSO control was *apo*-RXRα LBD containing 1% w/v DMSO. Data for the DMSO control and all other were obtained at a $v$ of 4 °C/min and in the presence of 1% DMSO.

**Figure 6.**
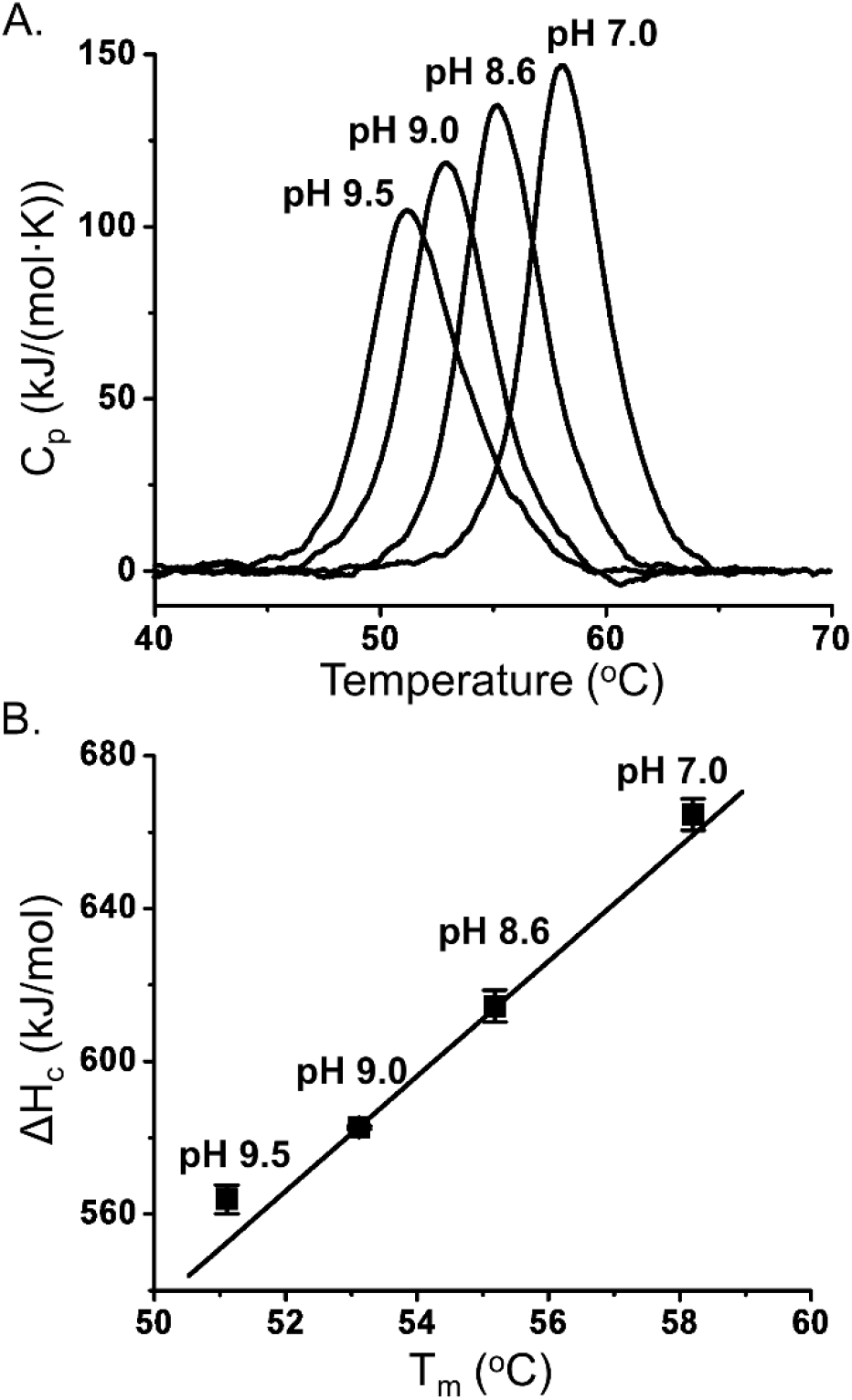
Determination of the unfolding heat capacity change (Δ*C_p_*^u^) of *apo*-RXRα LBD homodimers. A) DSC molar heat capacity profiles for 1.5 μM *apo*-RXR-LBD homodimer using a scan rate of 4 °C/min at different pH values (from left to right): 9.5, 9.0, 8.6, and 7.0. B) Δ*C_p_*^u^ was determined by a linear regression of Δ*H*_c_ versus *T*_m_. The R value for the linear fit was 0.99.

According to Harder *et al*. (42), the chemical denaturation of *apo*-RXRα LBD in guanidine-HCl is reversible and follows a three-state mechanism: N2 ↔ 2I ↔ 2U, wherein “I” is a monomeric, partially unfolded, intermediate. Although the Δ*G* change of the first step accounts for ~80% of the total unfolding Δ*G*, spectroscopy data support that the monomeric intermediate retains significant native secondary structure, and the two Trp residues remain partially buried. It is during the second step of unfolding where this unfolding intermediate, I, loses all secondary structures and the Trp are completely solvent exposed. The authors estimate that the Δ*G* of chemical denaturation for the first step is 35 ± 0.8 kJ/mol at 30 °C. From our study, the estimated thermal unfolding Δ*G* for *apo*-RXRα LBD homodimers at 30 °C was 39 ± 2 kJ/mol. Likewise, the spectral data on the thermally unfolded protein (Fig. 5) are similar to those estimated for the monomeric intermediate in guanidine-HCl. Guanidine-HCl is well known for its capability to prevent aggregation of partially or fully unfolded proteins because of its chaotropic and ionic properties (58–60). It is possible that, in our study, DSC detected the unfolding of the native dimer to a partially unfolded intermediate very similar to that detected by Harder *et al*. However, because of the lack of chaotropes such as Guanidine-HCl to stabilize a monomer, the unfolded state was thermally aggregated without a significant heat signature.

The ΔG of unfolding for *apo*-RXRα LBD was unusually low compared to other dimeric proteins, which may suggest a weak dimeric interface ready to dissociate into monomers at sub-micromolar concentrations to facilitate the formation of heterodimers with other receptors (42). However, our thermal unfolding data indicated that the dimer interface was relatively stable as it did not dissociate throughout the unfolding endotherm. This was more consistent with the DNA binding properties and transcriptional activities observed for the full-length RXR homodimer (3, 61). Moreover, the thermal unfolding of *apo*-RXRα LBD homodimer was accompanied by a relatively low unfolding Δ*H*_c_ of 673 kJ/mol at *T*_m_, i.e., a low specific heat of 12.5 J/g, which was substantially lower than the specific heat of a typical soluble globular protein at the same *T*_m_ (29 - 38 J/g according to Murphy and Freire (54)). The low specific heat may be caused by three factors. First, it may be due to incomplete unfolding as demonstrated by the CD and fluorescence spectroscopy data on the thermally unfolded protein. Second, it may be due to homodimers generally unfolding with lower specific heats than single domain proteins (62). Third, it may indicate that the native *apo*-homodimer is less compactly folded and more dynamic than typical, well-folded globular proteins. Such structural fluidity likely relates to the lack of rexinoid bound to its hydrophobic pocket and the dynamic helix 12 region of the domain (11, 12).

### II. Thermal Unfolding of *Holo*-RXRα LBD Homodimers Bound with Rexinoids

The stability of RXRα LBD bound with rexinoid (*holo*-RXRα LBD) was examined next to determine the relationship between rexinoid structure and *holo*-RXRα LBD unfolding energetics. Rexinoids UAB30, UAB110 and UAB111 quenched more than 90% of the protein fluorescence signal of *apo*-RXRα-LBD at 337 nm at 25 °C when the ratio of the protein to the rexinoids reached 1:1. The dissociation constants (*K*_d_) determined from the binding isotherm based on fluorescence quenching was 38 ± 14 nM for UAB30, 22 ± 6 nM for UAB110, and 2.4 ± 0.4 nM for UAB111.

DSF was first used to determine the rexinoid concentration required to saturate the binding sites at *T*_m_ because *K*_d_ was expected to be dependent on temperature. For UAB30, the DSF unfolding curve continuously shifted to higher *T*_m_ (**Fig. S4A**) when the ligand concentration was increased from 1.25 to 25 μM using a dimer protein concentration of 2.5 μM. There was no measurable increase in *T*_m_ when the rexinoid concentration was doubled to 50 μM, indicating that effective saturation was reached at rexinoid concentrations above 25 μM (5:1 molar ratio, based on one binding site/monomer). An increase in *T*_m_ of 4.4 ± 0.1 °C in the presence of UAB30 at a 2:1 rexinoid/monomer molar ratio was consistent with UAB30 strongly binding to the native homodimer at 25 °C and at elevated temperatures. For UAB110 and UAB111, the shifts in *T*_m_ in the presence of rexinoids at a 2:1 rexinoid/monomer molar ratio was 7 and 7.5 °C, respectively, which were substantially higher than the shifts caused by UAB30 (**Fig. S4B**), and consistent with their higher binding affinities.

To gather thermodynamic data on *holo*-RXRα LBD dimers, DSC was performed. The effect of DMSO, a cosolvent necessary for solubilizing the rexinoids, was first examined. Using a *v* of 4 °C/min, the *T*_m_, Δ*H*_c_ and Δ*H*_v_ values of *apo*-RXRα LBD homodimer determined by DSC in the presence of 1% w/v DMSO (Table 2) were within experimental errors of those values obtained in the absence of DMSO (Table 1). This sample was used as the point of reference (DMSO control) for comparison with *holo*-RXRα LBD dimers containing rexinoids. Rexinoid concentrations that caused maximum increases in *T*_m_ (30 μM for UAB30, and 10 μM for either UAB110 or UAB111) were used to minimize unwanted effects of the rexinoids at high concentrations (**Fig. S5**). To obtain extrapolated equilibrium parameters in the presence of rexinoids, DSC endotherms were measured at four different scan rates for the *holo*-RXRα LBD homodimer bound with UAB30 (RXRα LBD:UAB30, **Fig. S6A**). Similar to *apo*-RXRα LBD homodimer, only one unfolding transition was observed at a higher *T*_m_ due to the shift of unfolding equilibrium toward the native state caused by rexinoid binding to the *apo*-homodimer in the native state. Both *T*_m_ and Δ*H*_v_ displayed linear dependence on *v*^−1^, consistent with a Lumry-Eyring mechanism (**Fig. S6B, Table S1**). Similar to *apo*-RXRα LBD, Δ*H*_c_ values were independent of *v* (linear regression of Δ*H*_c_ against *v*^−1^ yielded a slope of zero). The extrapolated equilibrium unfolding parameters for the RXRα LBD:UAB30 *holo*-dimer were: *T*_m_^eq^ = 63.5 ± 0.1 °C, Δ*H*_v_^eq^ = 761 ± 38 kJ/mol, and Δ*H*_c_^eq^ = 798 ± 8 kJ/mol (using the average of the apparent Δ*H*_c_ at different *v* values). The Δ*H*_v_^eq^ and Δ*H*_c_^eq^ were within experimental error for the *holo*-homodimer, as found for the *apo*-homodimer. The scan rate dependence of *T*_m_ of RXRα LBD:UAB30 was also determined by DSF, and the results matched those determined by DSC (**Fig. S6B and Table S1**).

DSF was used to examine the effects of UAB110 and UAB111 on the stability of the homodimer. For *holo*-RXRα LBD homodimers in complex with either UAB110 or UAB111 (RXRα LBD:UAB110 and RXRα LBD:UAB111), the linear regression of *T*_m_ values of both complexes with respect to *v*^−1^ yielded slopes almost identical to that of *apo*-RXRα LBD or RXRα LBD:UAB30 homodimer (Fig. 7). This indicated the activation energy for the irreversible step in the Lumry-Eyring unfolding model was similar for both *holo*-homodimers and the *apo*-homodimer.

**Figure 7.**
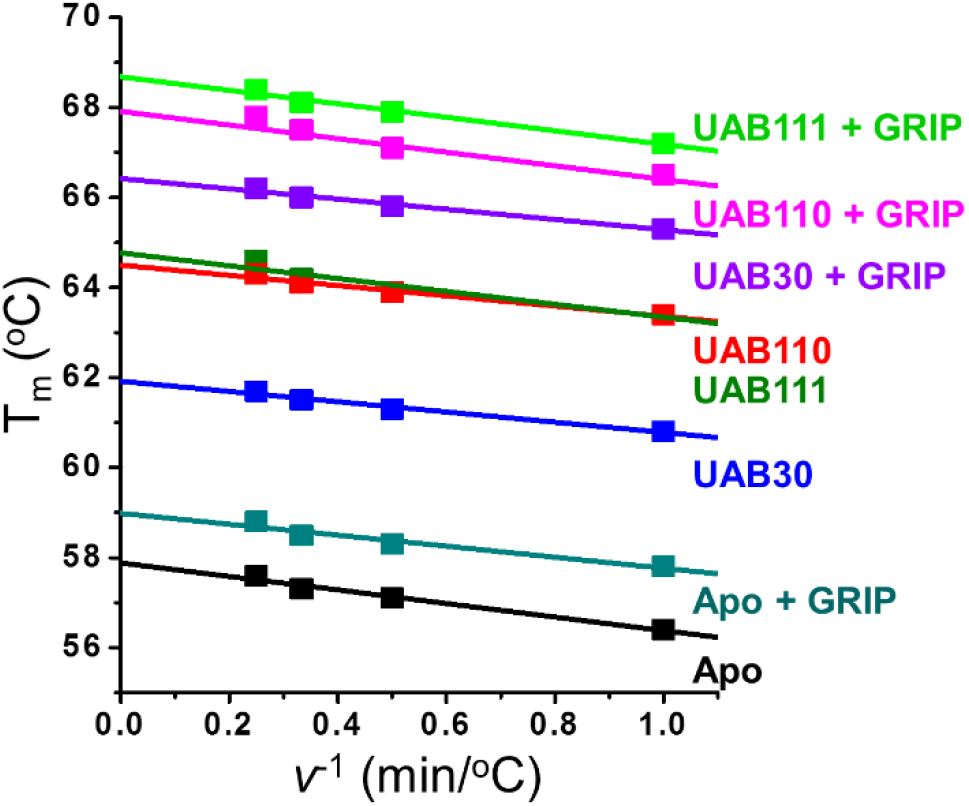
DSF *T*_m_ values as a function of *v*^−1^ for RXRα LBD homodimers with and without rexinoids or coactivator peptide. DSF *T*_m_ values of 2.5 μM RXRα LBD homodimer at pH 7.0, in the presence of no rexinoid or GRIP-1 (*Apo*), 0.4 mM GRIP-1 (Apo + GRIP), 30 μM UAB30 (UAB30), 30 μM UAB30 and 0.4 mM GRIP-1 (UAB30 + GRIP), 10 μM UAB110 (UAB110), 10 μM UAB110 and 0.4 mM GRIP-1 (UAB110 + GRIP), 10 μM UAB111 (UAB111), or 10 μM UAB111 and 0.4 mM GRIP-1 (UAB111 + GRIP). The scan rate, *v*, was 4.0 °C/min. The lines are linear regressions of *T*_m_ values with respect to *v*^−1^. All samples contained 1% DMSO.

To gather thermodynamic data, DSC measurements were conducted on RXRα LBD:UAB110 at a *v* of 4 °C/min. An unfolding *T*_m_ of 66.6 ± 0.5 °C was obtained, which was 3 °C higher than the *T*_m_ of RXRα LBD:UAB30 (Fig. 8 and Table 2). The Δ*H*_c_ was 849 ± 13 kJ/mol, which was approximately 260 kJ/mol higher than the *apo*-homodimer in 1% DMSO. RXRα LBD:UAB111 displayed similar unfolding parameters as UAB 110:RXRα LBD, with a *T*_m_ of 66.9 ± 0.3 °C and a Δ*H*_c_ of 840 ± 21 kJ/mol. Both RXRα LBD:UAB110 and RXRα LBD:UAB111 also exhibited a ΔHv /ΔHc ratio close to unity, consistent with the *holo*-RXRα LBD homodimers being the cooperative unfolding unit.

**Figure 8.**
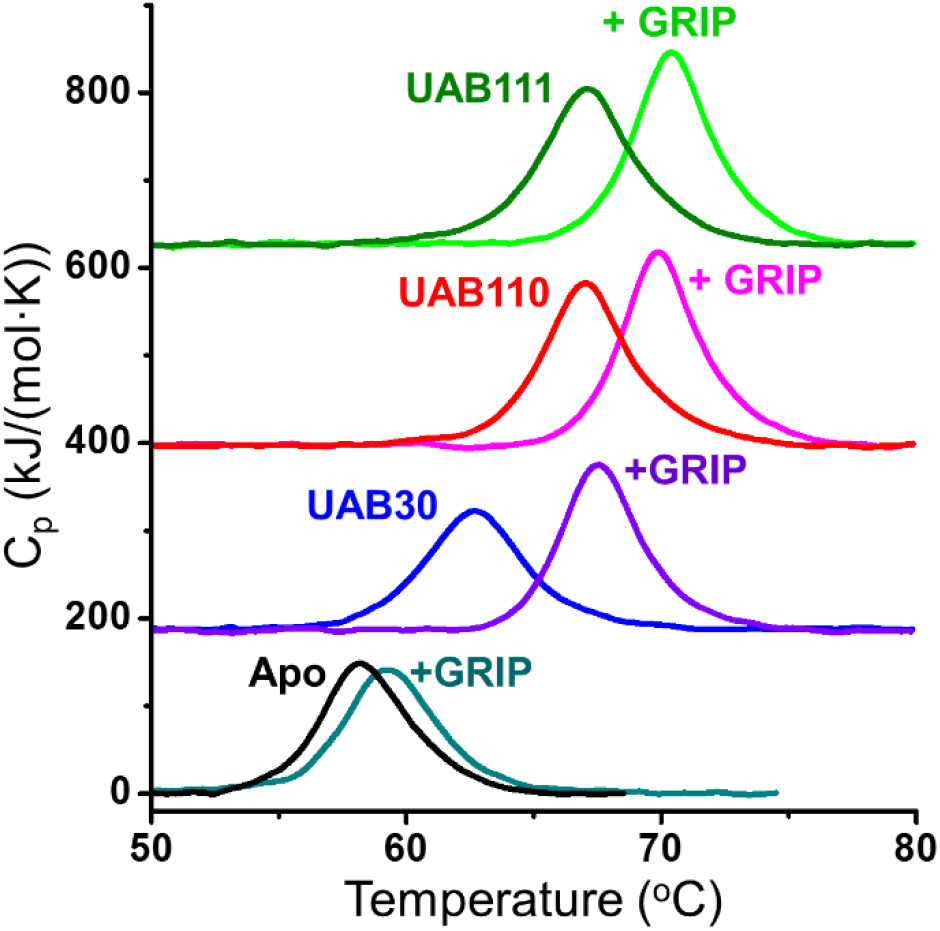
RXRα LBD is strongly stabilized by rexinoid binding and coactivator peptide GRIP-1. DSC molar heat capacity profiles of 1.5 μM RXRα LBD at pH 7.0, in the presence of no rexinoid or GRIP-1 (*Apo*), 0.4 mM GRIP-1 (+GRIP), 30 μM UAB30 (UAB30), 30 μM UAB30 and 0.4 mM GRIP-1 (UAB30 +GRIP), 10 μM UAB110 (UAB110), 10 μM UAB110 and 0.4 mM GRIP-1 (UAB110 +GRIP), 10 μM UAB111 (UAB111), or 10 μM UAB111 and 0.4 mM GRIP-1 (UAB111 +GRIP). The scan rate, *v*, was 4.0 °C/min. All samples contained 1% DMSO.

To determine the Δ*C_p_*^u^ values of the *holo*-RXRα LBD homodimers, *T*_m_ and Δ*H*_c_ were measured at pH 8.8 and 9.5. In a similar manner to *apo*-RXRα LBD homodimers, DSF was first used to evaluate whether pH modulates the degrees of stabilization caused by the presence of rexinoids (**Fig. S7** and **Table S2**). The shifts in *T*_m_ caused by rexinoid were within experimental errors at all three pH values. Using DSC, the measured *T*_m_ values and enthalpies were lower as pH decreased, but the relative positions and magnitudes of all DSC endotherm remained the same (**Fig. S8**). Δ*H*_c_ value of each *holo*-homodimer was fit linearly with respect to *T*_m_ to determine Δ*C_p_*^u^ values (Table 2).

Using Δ*C_p_*^u^, *T*_m_ and Δ*H*_c_ from DSC measurements at a *v* of 4 °C/min, Δ*H*, Δ*S*, and Δ*G* of unfolding at 37 °C for each *holo*-RXRα LBD homodimer were calculated (Table 2). Compared to the *apo*-RXRα LBD homodimer DMSO control, the *holo*-RXRα LBD homodimers had significantly higher Δ*G* values consistent with tight binding and strong stabilization of the native state by the rexinoids. The degrees that UAB110 and UAB111 stabilized the native *apo*-RXRα LBD homodimer (ΔΔ*G* = 19 and 20 kJ/mol, respectively) were notably higher than UAB30 (ΔΔ*G* = 12 kJ/mol), which is consistent with the improved binding affinities of UAB110 and UAB111 over that of UAB30.

The difference in Δ*H* between the DMSO control and *holo*-RXRα LBD homodimers (ΔΔ*H*) at 37 °C was essentially zero for each of the three rexinoids. A near-zero value for ΔΔ*H* suggested that the binding of rexinoid to *apo*-RXRα LBD at physiological temperatures was accompanied by a small binding enthalpy. In order to gather information of the thermodynamics of rexinoid binding to *apo*-RXRα LBD, ITC measurements were performed on UAB30 between 10 and 30 °C. No ITC isotherms were obtained due to low solubility of the rexinoid in water. When reverse titrations were performed at these temperatures, the measured heat changes were within the noise levels (Fig. S9), which suggested a low-enthalpy binding reaction at these temperatures, consistent with the DSC results.

The ligand binding pocket (LBP) of RXRα LBD containing UAB30 is lined with 16 hydrophobic residues contributed by 4 protein helices that surround the UAB rexinoids (11–12, 23–24). The main forces that stabilize the interactions between the rexinoids and LBD residues are the ionic interaction between the carboxylate group and Arg316, and numerous hydrophobic contacts throughout the binding pocket. Burying nonpolar atoms upon ligand binding leads to a negative heat capacity change (Δ*C_p_*^a^) (12, 38). Exposing these buried nonpolar surface areas upon protein unfolding and ligand dissociation increases the unfolding Δ*C_p_*^u^. The Δ*C_p_*^u^ of each of the three *holo*-RXRα LBD homodimers was slightly higher than that of the *apo*-RXRα LBD homodimer in 1% DMSO (+1 to 2 kJ/(mol·K)). The −*T*Δ*S* values of the *holo*-RXRα LBD homodimers were each less negative than that of the *apo*-RXRα LBD homodimer in DMSO (+17 to 31 kJ/mol), which contributed favorably to the Δ*G* values. Therefore, rexinoid binding to *apo*-RXRα LBD appeared to be entropically driven at physiological temperatures. Crystallographic data indicate that, upon binding to UAB110 or UAB111, the size of the LBP expanded nearly 20% (compared to LBP in the presence of UAB30) to accommodate the two larger rexinoids (Fig. 1D) (24). The contact surface areas of these two rexinoids were about 100 Å^2^ larger than those observed for UAB30. Burying more hydrophobic surfaces likely resulted in a more favorable entropy changes during the binding process, and hence the more favorable Δ*G* of folding for *holo*-RXRα LBD homodimers bound to UAB110 or UAB111. In addition, the increase in stability of the *holo*-RXRα LBD homodimers could also arise from reduced dynamics due to rexinoid binding. HDX MS studies have revealed that the dynamics of helices 3 and 11 are significantly decreased when UAB30 is bound to the homodimer (11), and reduced even further in the presence of UAB110 and UAB111 whose binding affinities were higher than that of UAB30 (unpublished data). In summary, the LBP was expanded but more thermodynamically stable when bound with UAB110 or UAB111. How these conformational changes affect coactivator binding was examined next.

### III. Thermal Unfolding of *Holo*-RXRα LBD Homodimers Bound with Rexinoids and a Coactivator Peptide

It was previously shown by using ITC measurements that the 13-mer coactivator peptide, GRIP-1 (Fig. 1C), binds to *holo*-RXRα LBD complexes with a micromolar *K*_d_ at 25 °C in an exothermic reaction with binding enthalpies in the range of −36 to −42 kJ/mol per monomer (11, 24). Because the binding affinity is expected to be weaker at *T*_m_, DSF was first used to determine the GRIP-1 concentration required to saturate the peptide binding site at *T*_m_ (**Fig. S10**). The *T*_m_ of UAB30:RXRα-LBD:GRIP-1 unfolding incrementally increased when GRIP-1 concentration was increased from 12.5 to 800 μM using a dimer protein concentration of 2.5 μM and 15 μM UAB30. A concentration of400 μM GRIP-1 was chosen for the collection of thermodynamic data on RXRα LBD homodimer bound with rexinoid and GRIP-1.

DSF was used to examine the dependence of *T*_m_ on *v* (Fig. 7). In the absence of rexinoids, GRIP-1 increased the *T*_m_ of *apo*-RXRα LBD by ~1 °C, consistent with its low affinity to the *apo*-homodimer at 25 °C (12). A much larger increase in *T*_m_ was observed in the presence of each rexinoid. All *T*_m_ values appeared linearly dependent on *v*^−1^, and the slopes of the linear fits were similar to those obtained in the absence of GRIP-1. This again suggested that the activation energies of the irreversible step of the Lumry-Eyring unfolding mechanism were similar in the presence or absence of rexinoids or GRIP-1. RXRα LBD:UAB111:GRIP-1 complex displayed the highest extrapolated *T*_m_ of 68.7 ± 0.1 °C, nearly 11 °C higher than that of the *apo*-homodimer. RXRα LBD:UAB110:GRIP-1 complex had an extrapolated *T*_m_ of 67.9 ± 0.2 °C, and RXRα LBD: UAB30:GRIP-1 complex had an extrapolated *T*_m_ of 66.4 ± 0.1 °C. Using DSC (Fig. 8 and Table 2), similar increases in *T*_m_ were observed for the ternary complexes. RXRα LBD:UAB111:GRIP-1 had the highest *T*_m_ of 70.4 ± 0.2 °C, followed by RXRα LBD:UAB110:GRIP-1 with a *T*_m_ of 69.9 ± 0.5 °C, and RXRα LBD:UAB30:GRIP-1 with a *T*_m_ of 67.5 ± 0.3 °C. The ternary complexes also unfolded with higher Δ*H*_c_ values that were 130 to 150 kJ/mol more than the Δ*H*_c_ values of their corresponding *holo*-RXRα LBD homodimers without GRIP-1 (Table 2).

To determine the Δ*C_p_*^u^ values of *holo*-RXRα LBD homodimers in complex with GRIP-1, *T*_m_ and Δ*H*_c_ values were determined by DSC at pH 8.8 and 9.5 (**Fig. S8**). Δ*H*_c_ values of the *holo*-RXRα LBD homodimers in the presence of GRIP-1 were fit to a linear dependence on *T*_m_. From the slopes of the lines, Δ*C_p_*^u^ values were determined and listed in Table 2. The unfolding parameters of *apo*-RXRα LBD homodimer in the presence of 400 μM GRIP-1 were similar to the *apo*-homodimer with only slightly increased *T*_m_ and Δ*H*_c_, and almost identical Δ*C_p_*^u^. This indicated that significant dissociation of GRIP-1 likely occurred before the *apo*-homodimer unfolded because of low binding affinity at *T*_m_. For two of the three RXRα LBD:rexinoid:GRIP-1 complexes, Δ*C_p_*^u^ increased with respect to their corresponding *holo*-homodimer without GRIP-1, while there was little observed change in Δ*C_p_*^u^ for RXRα LBD:UAB30:GRIP-1. The largest increase in Δ*C_p_*^u^ was 3 kJ/(mol·K) for RXRα LBD:UAB111:GRIP-1, which was within the experimental errors of Δ*C_p_*^u^ measurement. Therefore, an average Δ*C_p_*^u^ value of 18 ± 2 kJ/(mol·K) was calculated from the experimentally determined values from the Δ*H*_c_ versus *T*_m_ plots for the three ternary complexes.

Using the above average Δ*C_p_*^u^ value and the *T*_m_ and Δ*H*_c_ values from DSC measurements at 4 °C/min, Δ*H*, Δ*S*, and Δ*G* of unfolding at 37 °C for the *holo*-RXRα LBD homodimers bound with GRIP-1 peptide were calculated (Table 2). Binding of GRIP-1 increased Δ*G* compared to measurements obtained in the absence of the coactivator peptide: 13 ± 4 kJ/mol for RXRα LBD:UAB30, 14 ± 3 kJ/mol for RXRα LBD:UAB110, and 13 ± 3 kJ/mol for RXRα LBD:UAB111. Using equation (6) based on the shifts in *T*_m_, the estimated *K*_d_ of GRIP-1 binding at *T*_m_ was 22 μM for RXRα LBD:UAB30, 26 μM for RXRα LBD:UAB110, and 24 μM for RXRα LBD:UAB111.

In our previous analysis using ITC (11), the binding enthalpy of GRIP-1 to RXRα LBD:UAB30 was determined to be −44.6 ± 0.1 kJ/mol per monomer at 30 °C, with a binding heat capacity change (Δ*C_p_*^a^) of −1.5 ± 0.1 kJ/(mol·K). Using these values, binding enthalpy at 67.5 °C (*T*_m_ of the RXRα LBD:UAB30:GRIP-1 ternary complex) was calculated to be −100 kJ/mol per monomer or −200 kJ/mol per dimer, which was similar to the measured value of −129 ± 35 kJ/mol. Based on the van’t Hoff equation (equation [7]), using the binding enthalpy and Δ*C_p_*^a^ determined by ITC, the *K*_d_ at 25 °C was calculated to be 0.74 μM, in close agreement with the *K*_d_ determined by ITC at this temperature (11).

To obtain an accurate measurement of the Δ*C_p_*^a^ values for GRIP-1 binding to RXRα LBD:UAB110 or RXRα LBD:UAB111, ITC was performed between 10 and 30 °C. (**Table S3**). As observed for RXRα LBD:UAB30 complex, the binding stoichiometry was nearly 1:1 (coactivator peptide/ RXRα LBD monomer unit). The free energy change of GRIP-1 binding to RXRα LBD:rexinoid complexes was driven strongly by a large negative enthalpy change. Using the temperature dependence of the enthalpy change for binding, the Δ*C_p_*^a^ was determined to be 1.19 ± 0.01 kJ/(mol·K) for GRIP-1 binding to RXRα LBD:UAB110, and 1.11 ± 0.07 kJ/(mol·K) for GRIP-1 binding to RXRα LBD:UAB111 (per monomer unit). The *K*_d_ at 25 °C was calculated to be 1.5 μM for RXRα LBD:UAB110 and 1.8 μM for RXRα LBD:UAB110, also in close agreement with the *K*_d_ determined by ITC at this temperature (24).

For each of the three *holo*-RXRα LBD homodimers in complex with UAB rexinoid, there was a general trend of increased unfolding Δ*H* values when GRIP-1 was present, whereas the −*T*Δ*S* values remained unchanged. This indicated that the increase in Δ*G* upon GRIP-1 binding to the *holo*-RXRα LBD homodimers was enthalpically driven, consistent with GRIP-1 binding to the *holo*-RXRα LBD homodimers being an enthalpically favored process (11, 12, 24). For these ternary complexes, the specific heat of unfolding at *T*_m_ was approximately 18 J/g, a nearly 50% increase from the value of *apo*-RXRα LBD homodimer. While still less than the average specific heats of small, single-domain proteins, this increase is consistent with the ternary complexes with reduced structural fluidity and more compact folding in the presence of both rexinoid and coactivator peptide. Overall, the energetic changes induced by GRIP-1 binding to each of the three *holo*-RXRα LBD homodimers studied here appeared to be indistinguishable from one another, although each *holo*-homodimer displayed different stability that correlated to their structure and rexinoid-binding affinity. It is possible other coactivators may be more sensitive to these differences than GRIP-1. In this case, while there are clear differences in the binding properties of the three rexinoids, the result for the LBD as a whole is the formation of an AF-2 site that appears to be equally ready for GRIP-1 binding.

## CONCLUSIONS

It is well recognized that the functions of a nuclear receptor are relatively compartmentalized into its structural domains (1). Isolated DBD and LBD are well-structured and maintain their separate functions in vitro. The LBD is especially important because it harbors the ligand recognition site, the coregulator interaction site, and the main dimerization interface. In this study we have shown that *apo*-RXRα LBD homodimer, similar to other autonomously folded domains of multi-domain proteins, is favored in the folded state by nearly 33 kJ/mol in free energy change at 37 °C, driven by a favorable enthalpic change. On a per residue basis, the enthalpy value is almost 50% less than other small globular proteins, most likely due to the less compact nature of the *apo*-homodimer arising from the unfilled hydrophobic ligand binding pocket and a dynamic c-terminal end not interacting well with the core of the homodimer. Upon rexinoid binding, the hydrophobic pocket is filled, stabilized by numerous contacts between the rexinoid and nonpolar residues, along with the ionic interaction between the carboxylate group and a charged residue, and a hydrogen-bonding network with bound water molecules and hydrophilic groups (11, 24). These hydrophobic and hydrophilic interactions fill the interior and help bridge the helices interacting with the rexinoids (H3, H5, H7 and H11) to those at the terminal end where the coactivator peptide binds (H4 and H12). We clearly demonstrate for the first time that rexinoid binding to *apo*-RXRα LBD is entropically driven at physiological temperatures. The rexinoids enhance stability of the homodimer complex by 12 to 20 kJ/mol, depending on their structure. The ΔΔ*G* of stabilization is less for the smaller UAB30 and more for the larger and more hydrophobic UAB110 and UAB111. Even so, helix 12 does not reside in a helical structure interacting with H4 and H11 to form the coactivator binding site. Instead, it predominates in a less ordered, dynamic state (11). Only upon coactivator peptide binding does H12 join the rest of the structure and connects structurally to the compact homodimer domain (11, 24). This binding results in an additional stability to the homodimer complex of approximately 14 kJ/mol. In reported transcriptional assays (24), each rexinoid is a full agonist with different potencies. Structurally, the homodimer complexes with GRIP-1 are essentially identical with only very small changes in dynamics. It seems as if GRIP-1 does not sense the subtle differences in the LBP conformation caused by different rexinoids as long as the LBP is occupied and the GRIP-1 binding site is formed. Thus, the observed similarities in the reported crystal structures. This indicates that the difference in agonist potency may be the result of RXR heterodimer interactions rather than differential coactivator affinity. Alternatively, we may find that different RXR coactivator LXXLL motifs have unique binding characteristics with unique RXR ligands. These studies lay the groundwork for future thermodynamic studies of both RXR heterodimers and other coactivators.

## Supporting information

Yang et al. Supplemental

## ASSOCIATED CONTENT

### Supporting Information

Scan-rate dependence of DSF unfolding transitions of *apo*-RXRα LBD (Figure S1), Simulated DSC unfolding transitions of a hypothetical dimeric protein using two-state models with or without dimer dissociation (Figure S2), pH dependence of DSF unfolding transitions (Figure S3), Determination of saturating rexinoid concentrations at *T*_m_ (Figure S4), Nonspecific protein destabilization by UAB110 and UAB111 at high concentrations (Figure S5), Equilibrium unfolding parameters of RXRα LBD:UAB30 obtained by extrapolation to infinite scan-rate (Figure S6), DSC parameters of RXRα LBD:UAB30 at different scan-rates and extrapolated to infinite scan-rate (Table S1), DSF *T*_m_ of RXRα LBD bound with UAB30 at different pH values (Figure S7), DSF *T*_m_-shifts of *holo*-RXRα LBD do not vary with pH (Table S2), DSC of *apo*-RXRα LBD, *holo*-RXRα LBD with and without GRIP-1 at pH 8.8 and pH 9.5 (Figure S8), ITC of *apo*-RXRα LBD and UAB30 at different temperatures (Figure S9), DSF Tm of UAB30:RXRα LBD as a function of GRIP-1 concentration (Figure S10), Summary of ITC measurements of GRIP-1 to RXRα LBD:rexinoid complexes (Table S3)

### Accession Codes

The UniProt accession code for the retinoid X receptor-alpha is P19793.

### Funding Sources

NIH Grant P01 CA210946

## ACKNOWLEDGEMENT

The authors would like to thank Dr. Christie Brouillete for her helpful comments on the manuscript. This work was supported by NIH grant # P01 CA210946. Access to the VP-Capillary DSC, Auto-iTC200, and Prometheus NT.48 instruments was provided by the Biocalorimetry Lab supported by the NIH Shared Instrumentation Grant # 1S10RR026478 and Shared Facility Program of the UAB Comprehensive Cancer Center, Grant # 316851.

## ABBREVIATIONS

RXRα LBD: Retinoid X Receptor-alpha Ligand Binding Domain
LBP: ligand binding pocket
9*c*RA: 9-cis retinoic acid
GRIP-1: glucocorticoid receptor interacting protein-1
RXRα LBD:UAB30: RXRα LBD bound to UAB30
RXRα LBD:UAB30:GRIP-1: RXRα LBD bound to UAB30 and GRIP-1
RXRα LBD:UAB110: RXRα LBD bound to UAB110; RXRα
LBD:UAB110:GRIP-1: RXRα LBD bound to UAB110 and GRIP-1; RXRα
LBD:UAB111: RXRα LBD bound to UAB111; RXRα
LBD:UAB111:GRIP-1: RXRα LBD bound to UAB111 and GRIP-1
SRC: steroid receptor coactivator
DSC: differential scanning calorimetry
*v*: scan or heating rate
*T*_m_: thermal unfolding temperature
Δ*H*_v_: van’t Hoff enthalpy of unfolding
Δ*H*_c_: calorimetric enthalpy
Δ*G*: Gibbs free energy of unfolding
Δ*S*: entropy of unfolding
Δ*C_p_*^u^: unfolding heat capacity change
*K*_u_: unfolding equilibrium constant
DSF: differential scanning fluorimetry
F_350_/F_330_: ratio of fluorescence intensity at 350 nm divided by fluorescence intensity at 330 nm
λ_ex_: excitation wavelength for fluorescence measurements
CD: Circular Dichroism
ITC: isothermal titration calorimetry
Δ*H*_a_: binding enthalpy
Δ*C_p_*^a^: binding heat capacity change
*K*_a_: ligand binding constant
*K*_d_: ligand dissociation constant
Δ*C_p_*^a^: heat capacity change upon binding
HDX-MS: hydrogen-deuterium exchange mass spectrometry
N: native folded protein
U: reversibly unfolded protein
D: irreversibly denatured protein
I: partially unfolded intermediate
R: rexinoid
P: coactivator peptide GRIP-1

## Footnotes

1 The extent of unfolding at temperature *T* was determined by the ratio of integrated area from 35 °C to *T*, divided by the total area under the DSC curve.

## REFERENCES

1. Dawson, M. I., and Xia, Z. (2012). The retinoid X receptors and their ligands. Biochim Biophys Acta, 1821(1), 21–56.

2. Mangelsdorf, D. J., Thummel, C., Beato, M., Herrlich, P., Schutz, G., Umesono, K.,… Evans, R. M. (1995). The nuclear receptor superfamily: the second decade. Cell, 83(6), 835–839.

3. Zhao, Q., Chasse, S. A., Devarakonda, S., Sierk, M. L., Ahvazi, B., and Rastinejad, F. (2000). Structural basis of RXR-DNA interactions. J Mol Biol, 296(2), 509–520.

4. Bourguet, W., Ruff, M., Chambon, P., Gronemeyer, H., and Moras, D. (1995). Crystal structure of the ligand-binding domain of the human nuclear receptor RXR-alpha. Nature, 375(6530), 377–382.

5. Egea, P. F., Mitschler, A., Rochel, N., Ruff, M., Chambon, P., and Moras, D. (2000). Crystal structure of the human RXRalpha ligand-binding domain bound to its natural ligand: 9-cis retinoic acid. EMBO J, 19(11), 2592–2601.

6. Egea, P. F., Mitschler, A., and Moras, D. (2002). Molecular recognition of agonist ligands by RXRs. Mol Endocrinol, 16(5), 987–997.

7. Eberhardt, J., McEwen, A. G., Bourguet, W., Moras, D., and Dejaegere, A. (2019). A revisited version of the apo structure of the ligand-binding domain of the human nuclear receptor retinoic X receptor alpha. Acta Crystallogr F Struct Biol Commun, 75(Pt 2), 98–104.

8. Yan, X., Broderick, D., Leid, M. E., Schimerlik, M. I., and Deinzer, M. L. (2004). Dynamics and ligand-induced solvent accessibility changes in human retinoid X receptor homodimer determined by hydrogen deuterium exchange and mass spectrometry. Biochemistry, 43(4), 909–917.

9. Chalmers, M. J., Busby, S. A., Pascal, B. D., He, Y., Hendrickson, C. L., Marshall, A. G., and Griffin, P. R. (2006). Probing protein ligand interactions by automated hydrogen/deuterium exchange mass spectrometry. Anal Chem, 78(4), 1005–1014.

10. Lu, J., Cistola, D. P., and Li, E. (2006). Analysis of ligand binding and protein dynamics of human retinoid X receptor alpha ligand-binding domain by nuclear magnetic resonance. Biochemistry, 45(6), 1629–1639.

11. Boerma, L. J., Xia, G., Qui, C., Cox, B. D., Chalmers, M. J., Smith, C. D., Lobo-Ruppert, S., Griffin, P. R., Muccio, D. D., and Renfrow, M. B. (2014). Defining the communication between agonist and coactivator binding in the retinoid X receptor alpha ligand binding domain. J Biol Chem, 289(2), 814–826.

12. Xia, G., Boerma, L. J., Cox, B. D., Qiu, C., Kang, S., Smith, C. D., Renfrow, M. B., and Muccio, D. D. (2011). Structure, energetics, and dynamics of binding coactivator peptide to the human retinoid X receptor alpha ligand binding domain complex with 9-cis-retinoic acid. Biochemistry, 50(1), 93–105.

13. Chandra, V., Huang, P., Hamuro, Y., Raghuram, S., Wang, Y., Burris, T. P., and Rastinejad, F. (2008). Structure of the intact PPAR-gamma-RXR-nuclear receptor complex on DNA. Nature, 456(7220), 350–356.

14. Chandra, V., Wu, D., Li, S., Potluri, N., Kim, Y., and Rastinejad, F. (2017). The quaternary architecture of RARbeta-RXRalpha heterodimer facilitates domain-domain signal transmission. Nat Commun, 8(1), 868.

15. Zhang, J., Chalmers, M. J., Stayrook, K. R., Burris, L. L., Garcia-Ordonez, R. D., Pascal, B. D., Burris T. P., Dodge J. A., and Griffin, P. R. (2010). Hydrogen/deuterium exchange reveals distinct agonist/partial agonist receptor dynamics within vitamin D receptor/retinoid X receptor heterodimer. Structure, 18(10), 1332–1341.

16. Lou, X., Toresson, G., Benod, C., Suh, J. H., Philips, K. J., Webb, P., and Gustafsson, J. A. (2014). Structure of the retinoid X receptor alpha-liver X receptor beta (RXRalpha-LXRbeta) heterodimer on DNA. Nat Struct Mol Biol, 21(3), 277–281.

17. Kojetin, D. J., Matta-Camacho, E., Hughes, T. S., Srinivasan, S., Nwachukwu, J. C., Cavett, V., Nowak J., Chalmers, M. J., Marciano D. P., Kamenecka, T. M., Shulman, A. L., Rance, M., Griffin, P. R., Bruning, J. B., and Nettles, K. W. (2015). Structural mechanism for signal transduction in RXR nuclear receptor heterodimers. Nat Commun, 6, 8013.

18. Heery, D. M., Kalkhoven, E., Hoare, S., and Parker, M. G. (1997). A signature motif in transcriptional co-activators mediates binding to nuclear receptors. Nature, 387(6634), 733–736.

19. McKenna, N. J., and O’Malley, B. W. (2002). Combinatorial control of gene expression by nuclear receptors and coregulators. Cell, 108(4), 465–474.

20. Nahoum, V., Perez, E., Germain, P., Rodriguez-Barrios, F., Manzo, F., Kammerer, S.,… Bourguet, W. (2007). Modulators of the structural dynamics of the retinoid X receptor to reveal receptor function. Proc Natl Acad Sci U S A, 104(44), 17323–17328.

21. Zhang, H., Li, L., Chen, L., Hu, L., Jiang, H., and Shen, X. (2011). Structure basis of bigelovin as a selective RXR agonist with a distinct binding mode. J Mol Biol, 407(1), 13–20.

22. Lippert, W. P., Burschka, C., Gotz, K., Kaupp, M., Ivanova, D., Gaudon, C.,… Tacke, R. (2009). Silicon analogues of the RXR-selective retinoid agonist SR11237 (BMS649): chemistry and biology. ChemMedChem, 4(7), 1143–1152.

23. Atigadda, V. R., Xia, G., Desphande, A., Boerma, L. J., Lobo-Ruppert, S., Grubbs, C. J.,… Muccio, D. D. (2014). Methyl substitution of a rexinoid agonist improves potency and reveals site of lipid toxicity. J Med Chem, 57(12), 5370–5380.

24. Atigadda, V. R., Xia, G., Deshpande, A., Wu, L., Kedishvili, N., Smith, C. D., Krontiras, H., Bland, K. I., Grubbs, C. J., Brouillette, W. J., and Muccio, D. D. (2015). Conformationally Defined Rexinoids and Their Efficacy in the Prevention of Mammary Cancers. J Med Chem, 58(19), 7763–7774.

25. Grubbs, C. J., Lubet, R. A., Atigadda, V. R., Christov, K., Deshpande, A. M., Tirmal, V., Xia, G., Bland, K. I., Eto, I., Brouillette, W. J., and Muccio, D. D. (2006). Efficacy of new retinoids in the prevention of mammary cancers and correlations with short-term biomarkers. Carcinogenesis, 27(6), 1232–1239.

26. Muccio, D. D., Atigadda, V. R., Brouillette, W. J., Bland, K. I., Krontiras, H., and Grubbs, C. J. (2017). Translation of a Tissue-Selective Rexinoid, UAB30, to the Clinic for Breast Cancer Prevention. Curr Top Med Chem, 17(6), 676–695.

27. Garner, E. F., Stafman, L. L., Williams, A. P., Aye, J. M., Goolsby, C., Atigadda, V. R., Moore, B. P., Nan, L., Stewart, J. E., Hjelmeland, A. B., Friedman, G. K., and Beierle, E. A. (2018). UAB30, a novel RXR agonist, decreases tumorigenesis and leptomeningeal disease in group 3 medulloblastoma patient-derived xenografts. J Neurooncol, 140(2), 209–224.

28. Chou, C. F., Hsieh, Y. H., Grubbs, C. J., Atigadda, V. R., Mobley, J. A., Dummer, R., Muccio, D.D., Eto, I., Elmets C. A., Garvet, W. T., and Chang, P. L. (2018). The retinoid X receptor agonist, 9-cis UAB30, inhibits cutaneous T-cell lymphoma proliferation through the SKP2-p27kip1 axis. J Dermatol Sci, 90(3), 343–356.

29. Gniadecki, R., Assaf, C., Bagot, M., Dummer, R., Duvic, M., Knobler, R., Ranki, A., Schwandt, P., and Whittaker, S. (2007). The optimal use of bexarotene in cutaneous T-cell lymphoma. Br J Dermatol, 157(3), 433–440.

30. Muccio, D. D., Brouillette, W. J., Breitman, T. R., Taimi, M., Emanuel, P. D., Zhang, X., Chen, G., Sani, B. P., Venepally, P., Reddy, L., Alam, M., Simpson-Herren, L., and Hill, D. L. (1998). Conformationally defined retinoic acid analogues. 4. Potential new agents for acute promyelocytic and juvenile myelomonocytic leukemias. J Med Chem, 41(10), 1679–1687.

31. Atigadda, V. R., Vines, K. K., Grubbs, C. J., Hill, D. L., Beenken, S. L., Bland, K. I., Brouillette, W. J., and Muccio, D. D. (2003). Conformationally defined retinoic acid analogues. 5. Large-scale synthesis and mammary cancer chemopreventive activity for (2E,4E,6Z,8E)-8-(3’,4’-dihydro-1’(2’H)-naphthalen-1’-ylidene)-3,7-dimethyl-2,4,6-octatrienoic acid (9cUAB30). J Med Chem, 46(17), 3766–3769.

32. Kolesar, J. M., Hoel, R., Pomplun, M., Havighurst, T., Stublaski, J., Wollmer, B., Krontiras, H., Brouillette, W. J., Muccio, D. D., Kim, K., Grubbs, C. J., and Bailey, H. E. (2010). A pilot, first-in-human, pharmacokinetic study of 9cUAB30 in healthy volunteers. Cancer Prev Res (Phila), 3(12), 1565–1570.

33. Kolesar, J. M., Andrews, S., Green, H., Havighurst, T. C., Wollmer, B. W., DeShong, K., Laux, D. E., Krontiras, H., Muccio, D. D., Kim, K., Grubbs, C. J., House, M. G., Parnes, H. L., Heckman-Stoddard, B. M., and Bailey, H. H. (2019). A Randomized, Placebo-Controlled, Double-Blind, Dose Escalation, Single Dose, and Steady-State Pharmacokinetic Study of 9cUAB30 in Healthy Volunteers. Cancer Prev Res (Phila), 12(12), 903–912.

34. Desphande, A., Xia, G., Boerma, L. J., Vines, K. K., Atigadda, V. R., Lobo-Ruppert, S., Grubbs, C. J., Moeinpour, F. L., Smith, C. D., Christov, K, Brouillette W. J., and Muccio, D. D. (2014). Methyl-substituted conformationally constrained rexinoid agonists for the retinoid X receptors demonstrate improved efficacy for cancer therapy and prevention. Bioorg Med Chem, 22(1), 178–185.

35. Doyle, M. L. (1997). Characterization of binding interactions by isothermal titration calorimetry. Curr Opin Biotechnol, 8(1), 31–35.

36. Ward, W. H., and Holdgate, G. A. (2001). Isothermal titration calorimetry in drug discovery. Prog Med Chem, 38, 309–376.

37. Brandts, J. F., and Lin, L. N. (1990). Study of strong to ultratight protein interactions using differential scanning calorimetry. Biochemistry, 29(29), 6927–6940.

38. Waldron, T. T., and Murphy, K. P. (2003). Stabilization of proteins by ligand binding: application to drug screening and determination of unfolding energetics. Biochemistry, 42(17), 5058–5064.

39. Schimerlik, M. I., Peterson, V. J., Hobbs, P. D., Dawson, M. I., and Leid, M. (1999). Kinetic and thermodynamic analysis of 9-cis-retinoic acid binding to retinoid X receptor alpha. Biochemistry, 38(21), 6732–6740.

40. Yang, Z., and Brouillette, C. G. (2016). A Guide to Differential Scanning Calorimetry of Membrane and Soluble Proteins in Detergents. Methods Enzymol, 567, 319–358.

41. Pantoliano, M. W., Petrella, E. C., Kwasnoski, J. D., Lobanov, V. S., Myslik, J., Graf, E., Carver, T., Asel, E., Springer, B. A., Lane, P., and Salemme, F. R. (2001). High-density miniaturized thermal shift assays as a general strategy for drug discovery. J Biomol Screen, 6(6), 429–440.

42. Harder, M. E., Malencik, D. A., Yan, X., Broderick, D., Leid, M., Anderson, S. R., Deinzer, M. L., and Schimerlik, M. I. (2009). Equilibrium unfolding of the retinoid X receptor ligand binding domain and characterization of an unfolding intermediate. Biophys Chem, 141(1), 1–10.

43. Privalov, P. L. (1979). Stability of proteins: small globular proteins. Adv Protein Chem, 33, 167–241.

44. Garidel, P., Hegyi, M., Bassarab, S., and Weichel, M. (2008). A rapid, sensitive and economical assessment of monoclonal antibody conformational stability by intrinsic tryptophan fluorescence spectroscopy. Biotechnol J, 3(9-10), 1201–1211.

45. Zelent, B., Sharp, K. A., and Vanderkooi, J. M. (2010). Differential scanning calorimetry and fluorescence study of lactoperoxidase as a function of guanidinium-HCl, urea, and pH. Biochim Biophys Acta, 1804(7), 1508–1515.

46. Yin, S. W., Tang, C. H., Yang, X. Q., and Wen, Q. B. (2011). Conformational study of red kidney bean (Phaseolus vulgaris L.) protein isolate (KPI) by tryptophan fluorescence and differential scanning calorimetry. J Agric Food Chem, 59(1), 241–248.

47. Thiagarajan, G., Semple, A., James, J. K., Cheung, J. K., and Shameem, M. (2016). A comparison of biophysical characterization techniques in predicting monoclonal antibody stability. MAbs, 8(6), 1088–1097.

48. Freire, E., van Osdol, W. W., Mayorga, O. L., and Sanchez-Ruiz, J. M. (1990). Calorimetrically determined dynamics of complex unfolding transitions in proteins. Annu Rev Biophys Biophys Chem, 19, 159–188.

49. Senisterra, G. A., Markin, E., Yamazaki, K., Hui, R., Vedadi, M., and Awrey, D. E. (2006). Screening for ligands using a generic and high-throughput light-scattering-based assay. J Biomol Screen, 11(8), 940–948.

50. Lumry, R., and Eyring, H. (1954). Conformation Changes of Proteins. The Journal of Physical Chemistry, 58(2), 110–120.

51. Sanchez-Ruiz, J. M. (1992). Theoretical analysis of Lumry-Eyring models in differential scanning calorimetry. Biophys J, 61(4), 921–935.

52. Tello-Solis, S. R., and Hernandez-Arana, A. (1995). Effect of irreversibility on the thermodynamic characterization of the thermal denaturation of Aspergillus saitoi acid proteinase. Biochem J, 311 (Pt 3), 969–974.

53. Yang, Z. W., Tendian, S. W., Carson, W. M., Brouillette, W. J., Delucas, L. J., and Brouillette, C. G. (2004). Dimethyl sulfoxide at 2.5% (v/v) alters the structural cooperativity and unfolding mechanism of dimeric bacterial NAD+ synthetase. Protein Sci, 13(3), 830–841.

54. Murphy, K. P., and Freire, E. (1992). Thermodynamics of structural stability and cooperative folding behavior in proteins. Adv Protein Chem, 43, 313–361.

55. Nevzorov I, Redwood C, and Levitsky D. (2008) Stability of two beta-tropomyosin isoforms: effects of mutation Arg91Gly. J. Muscle Res. Cell Motil. 29 (6–8): 173–176.

56. Yang, Z., Zhou, Q., Mok, L., Singh, A., Swartz, D. J., Urbatsch, I. L., and Brouillette, C. G. (2017). Interactions and cooperativity between P-glycoprotein structural domains determined by thermal unfolding provides insights into its solution structure and function. Biochim Biophys Acta Biomembr, 1859(1), 48–60.

57. Robbins, R. J., Fleming, G. R., Beddard, G. S., Robinson, G. W., Thistlethwaite, P. J., and Woolfe, G. J. (1980). Photophysics of aqueous tryptophan: pH and temperature effects. Journal of the American Chemical Society, 102(20), 6271–6279.

58. Watlaufer, D. B., Malik, S. K., Stoller, L., and Coffin, R. L. (1964). Nonpolar Group Participation in the Denaturation of Proteins by Urea and Guanidinium Salts. Model Compound Studies. Journal of the American Chemical Society, 86(3), 508–514.

59. Nozaki, Y., and Tanford, C. (1970). The solubility of amino acids, diglycine, and triglycine in aqueous guanidine hydrochloride solutions. J Biol Chem, 245(7), 1648–1652.

60. Venkatesu, P., Lee, M. J., and Lin, H. M. (2007). Thermodynamic characterization of the osmolyte effect on protein stability and the effect of GdnHCl on the protein denatured state. J Phys Chem B, 111(30), 9045–9056.

61. Heery, D. M., Pierrat, B., Gronemeyer, H., Chambon, P., and Losson, R. (1994). Homo- and heterodimers of the retinoid X receptor (RXR) activated transcription in yeast. Nucleic Acids Res, 22(5), 726–731.

62. Privalov, P. L. (1982). Stability of proteins. Proteins which do not present a single cooperative system. Adv Protein Chem, 35, 1–104.

