## Supplementary material for "Stability of the Retinoid X Receptor-alpha Homodimer in the Presence and Absence of Rexinoid and Coactivator Peptide": Yang et al. Supplemental

**Figure S1. Scan-rate dependence of DSF unfolding transitions of *apo*-RXRα LBD**

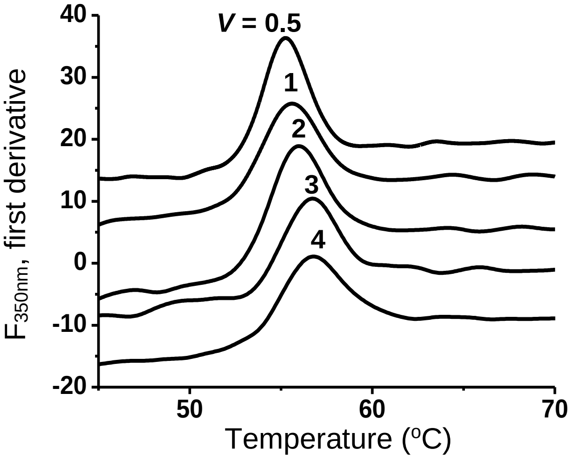


First derivative of the intrinsic fluorescence intensity of 2.5 μM *apo*-RXRα LBD homodimer at 350 nm ($F_{350nm}$) as a function of temperature at different scan rates (*v*): 0.5, 1.0, 2.0, 3.0, and 4.0 °C/min. Curves are Y-shifted for clarity. The excitation wavelength ($\lambda_{ex}$) was 290nm.

**Figure S2. Simulated DSC unfolding transitions of a hypothetical dimeric protein using two-state models with or without dimer dissociation**

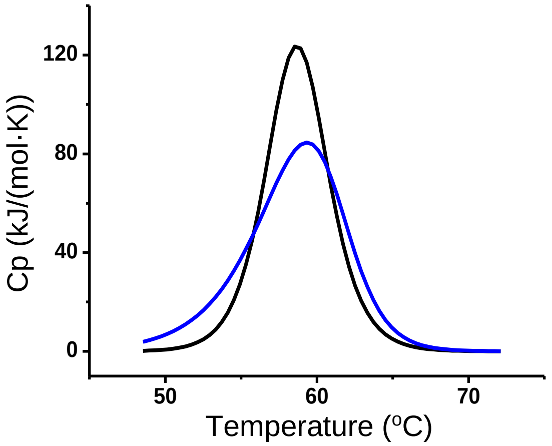


Black line: using the N ↔ U model (Model I) with $T_m$ = 58.7 $^o$C, and $\Delta H_c$ = 673 kJ/mol. Blue line: using the $N_2$ ↔ 2U model (Model II) with $T_m$ = 58.7 $^o$C, and $\Delta H_c$ = 673 kJ/mol. Note that the definition of $T_m$ for Model I is that the unfolding equilibrium constant, $K_u$, at $T_m$ is 1; whereas for Model II, the $K_u$ at $T_m$ is $2P_t$ where $P_t$ is the total protein concentration in terms of the dimer. When the blue curve was fitted using a non-dissociative model, the apparent $\Delta H_v/\Delta H_c$ was 0.69.

**Figure S3. pH dependence of DSF unfolding transitions**

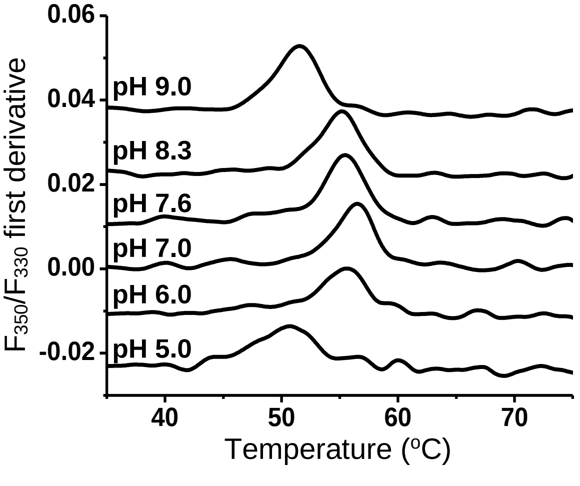


First derivative of the fluorescence ratio ($F_{350nm}/F_{330nm}$) of 1.5 μM *apo*-RXRα LBD homodimer as a function of temperature at different pH values: 9.0, 8.3, 7.6, 7.0, 6.0, and 5.0. The scan rate was 4 $^o$C/min. Curves are Y-shifted for clarity.

**Figure S4. Determination of saturating rexinoid concentrations at $T_m$**

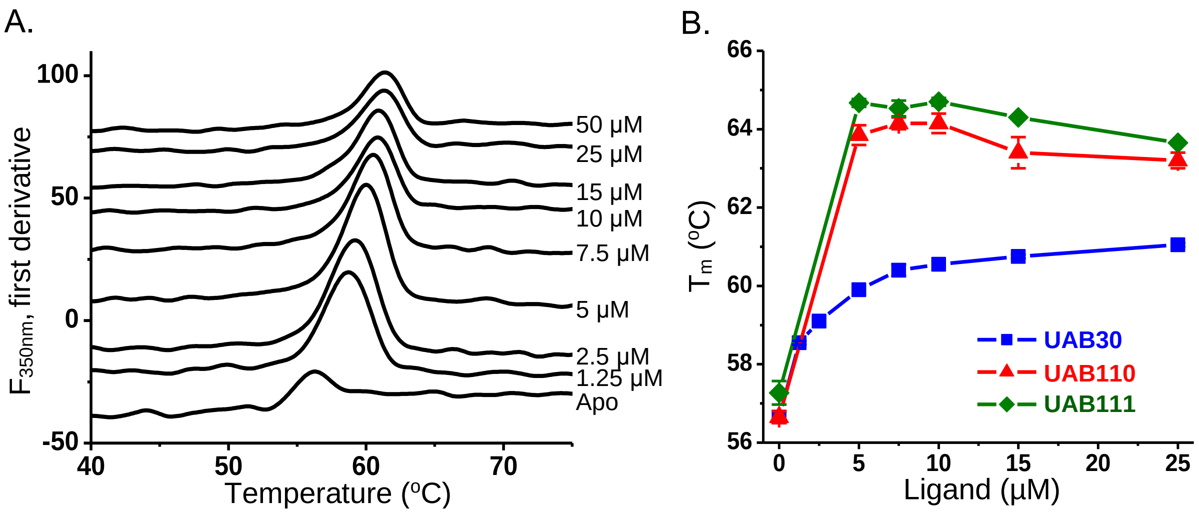


A) First derivative of the intrinsic fluorescence intensity of 2.5 μM RXRα LBD homodimer at 350 nm ($F_{350nm}$) as a function of temperature, in the presence of increasing concentrations of UAB30 from 0 to 50 μM. Curves are Y-shifted for clarity. B) DSF $T_m$ of 2.5 μM RXRα LBD homodimer in the presence of increasing rexinoids, determined from DSF curves as those shown in panel A.

For UAB110 and UAB111, as rexinoid concentration increased beyond 2:1 ligand/monomer molar ratio, the $T_m$ decreased. In addition, the magnitude of the fluorescence unfolding peak diminished significantly. No unfolding transition was observable by DSF at 50 μM (**Fig. S5A, see below**). To explore if the loss of the fluorescence unfolding signal was caused by quenching of Trp fluorescence by rexinoids, which absorb in the 320-350 nm range [Atigadda et al., 2003], the effect of rexinoids on the unfolding of a control protein, *B. subtilis* NAD synthetase (NADS), was investigated (**Fig. S5B**). NADS is a 60-kDa homodimeric enzyme, of which thermal unfolding mechanism and substrate binding properties have been determined previously [Yang et al., 2004]. The DSF curves of NADS did not shift to higher $T_m$ in the presence of rexinoids, indication of no specific interaction between this protein and the rexinoids. However, similar decreases in both NADS $T_m$ and the magnitude of DSF unfolding peak were observed with higher concentrations of UAB110 or UAB111, as well as the loss of unfolding signal at 50 μM. Additionally, light scattering of the DSF samples during unfolding indicated precipitation of UAB110 and UAB111 at concentrations above 15 μM (Fig. **S5D**), whereas UAB30 was soluble up to 50 μM (Fig. **S5C**). Therefore, the destabilizing effects at high rexinoid concentrations were nonspecific and possibly linked to their low water-solubility.

**Figure S5. Nonspecific protein destabilization by UAB110 and UAB111 at high concentrations**

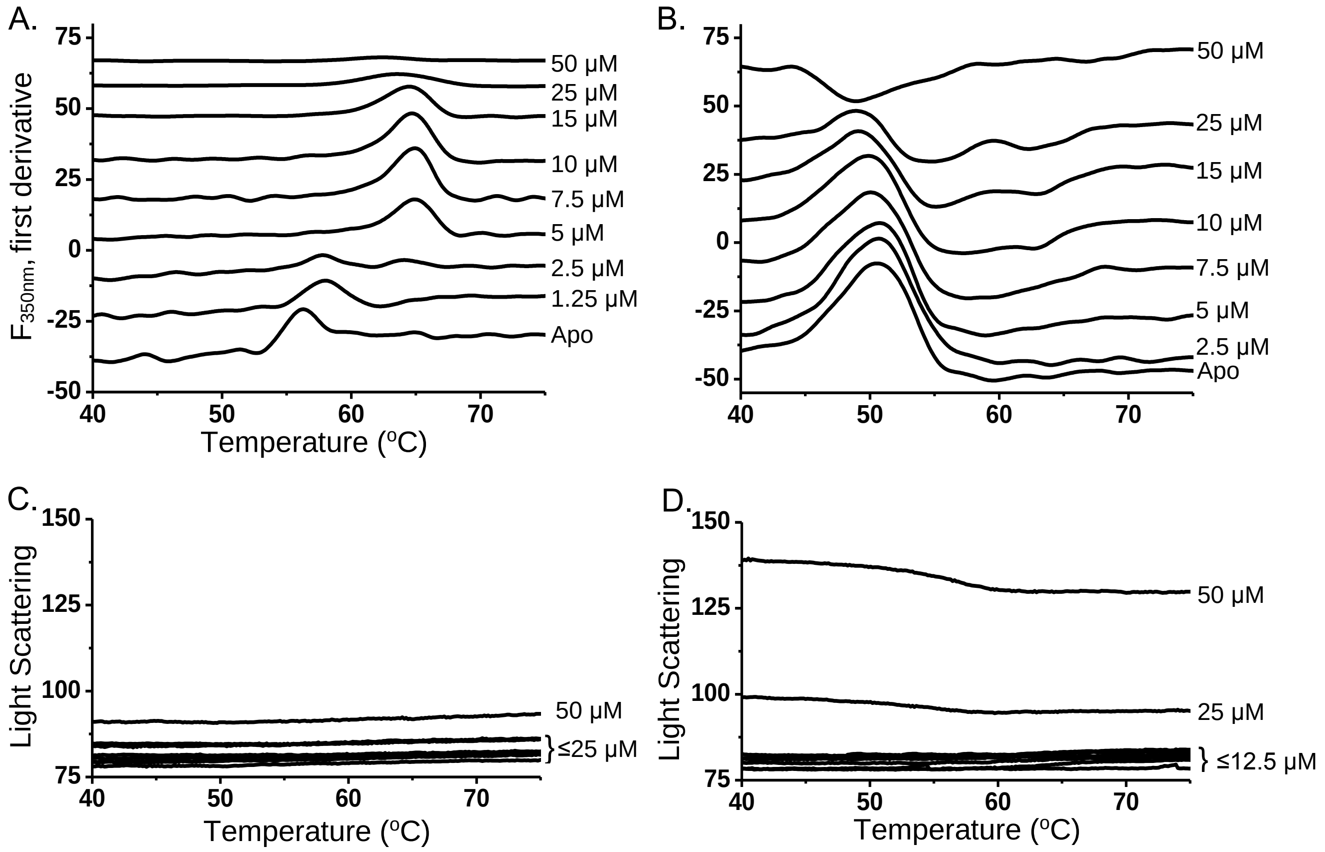


A) First derivative of the intrinsic fluorescence intensity of 2.5 µM RXRα LBD homodimer at 350 nm as a function of temperature in the presence of increasing concentrations of UAB111 from 0 to 50 µM. Curves are Y-shifted for clarity. B) First derivative of the intrinsic fluorescence intensity of 2.5 µM NAD synthetase homodimer (used as a control protein) at 350 nm as a function of temperature in the presence of increasing concentrations of UAB111 from 0 to 50 µM. Curves are Y-shifted for clarity. C) Light scattering intensity of the UAB30 samples in Fig. S4A recorded during DSF experiments. D) Light scattering intensity of the UAB111 samples in Fig. S5A recorded during DSF experiments.

**Figure S6. Equilibrium unfolding parameters of RXRα LBD:UAB30 obtained by extrapolation to infinite scan-rate**

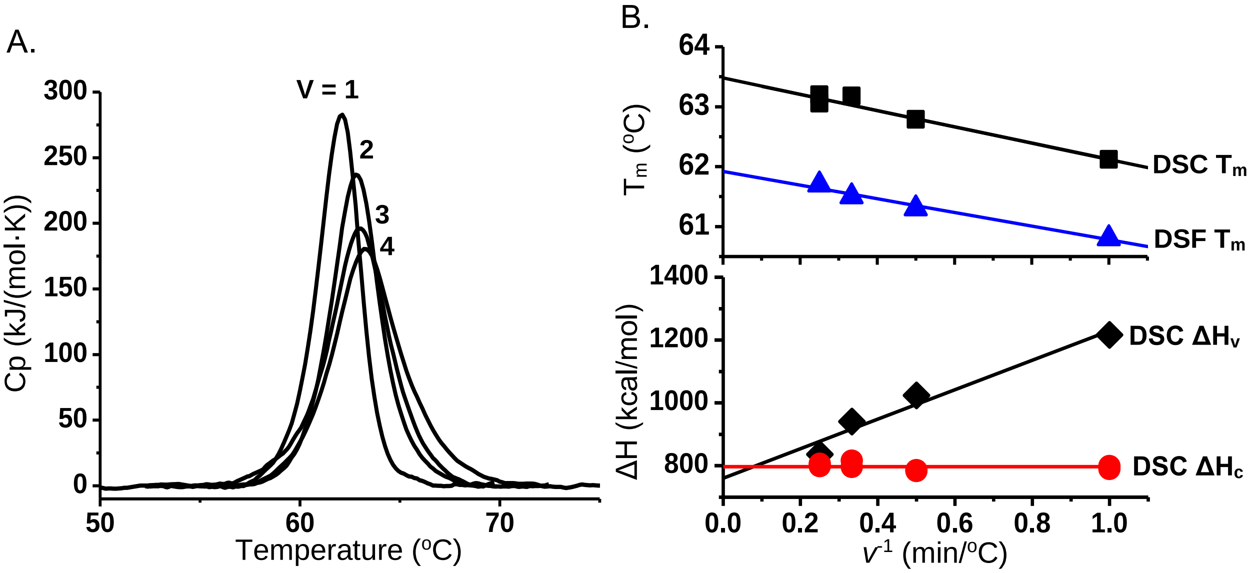


A) DSC molar heat capacity profiles for 1.5 μM RXRα LBD dimer in the presence of 30 μM UAB30 at different scan rates (*v*): 1.0, 2.0, 3.0, and 4.0 °C/min. B) $T_m$, $\Delta H_c$, and $\Delta H_v$ as a function of $v^{-1}$. Extrapolation to $v^{-1}$ = 0 yielded the equilibrium unfolding parameters (see Table S1). The red line is the average of the $\Delta H_c$'s at different *v*'s.

**Table S1. DSC parameters of RXRα LBD:UAB30 at different scan-rates and extrapolated to infinite scan-rate**

| Scan Rate | DSC | | | DSF |
|---|---|---|---|---|
| *v* (°C/min) | $T_m$ (°C) | $\Delta H_c$ (kJ/mol) | $\Delta H_v$ (kJ/mol) | $T_m$ (°C) |
| 1 | 62.1 ± 0.1 | 794 ± 4 | 1212 ± 4 | 60.80 ± 0.05 |
| 2 | 62.7 ± 0.1 | 786 ± 4 | 1028 ± 4 | 61.3 ± 0.1 |
| 3 | 63.0 ± 0.1 | 807 ± 13 | 945 ± 4 | 61.5 ± 0.1 |
| 4 | 63.1 ± 0.1 | 803 ± 4 | 836 ± 17 | 61.7 ± 0.2 |
| Infinite | 63.5 ± 0.1 | 798 ± 8 | 761 ± 38 | 61.9 ± 0.1 |

**Figure S7. DSF $T_m$ of RXRα LBD bound with UAB30 at different pH values**

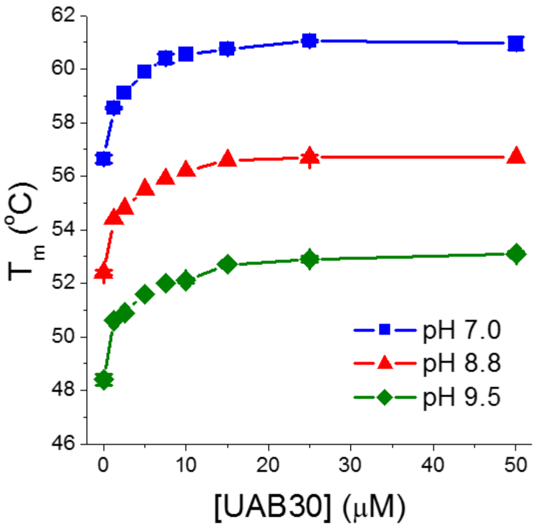


DSF $T_m$ of 2.5μM RXRα LBD homodimer in the presence of increasing UAB30 at three different pH values, determined from the first derivatives of fluorescence at 350 nm as a function of temperature. Buffer conditions were: 10mM sodium phosphate, pH 7.5, or 10mM sodium borate, pH 8.8, or 10 mM sodium borate, pH 9.5, with 50 mM NaCl, 2 mM EDTA, and 1 mM TCEP.

**Table S2. DSF $T_m$-shifts of *holo*-RXRα LBD at different pH values**

| | DSC $T_m$ shift (°C) | | |
|---|---|---|---|
| pH | 25 µM UAB30 | 10 µM UAB110 | 10 µM UAB111 |
| 7.0 | 4.5 ± 0.2 | 7.6 ± 0.4 | 8.1 ± 0.3 |
| 8.8 | 4.3 ± 0.2 | 7.8 ± 0.2 | 8.1 ± 0.2 |
| 9.5 | 4.5 ± 0.3 | 7.5 ± 0.4 | 8.0 ± 0.4 |

**Figure S8. DSC of *apo*-RXRα LBD, *holo*-RXRα LBD with and without GRIP-1 at pH 8.8 and pH 9.5**

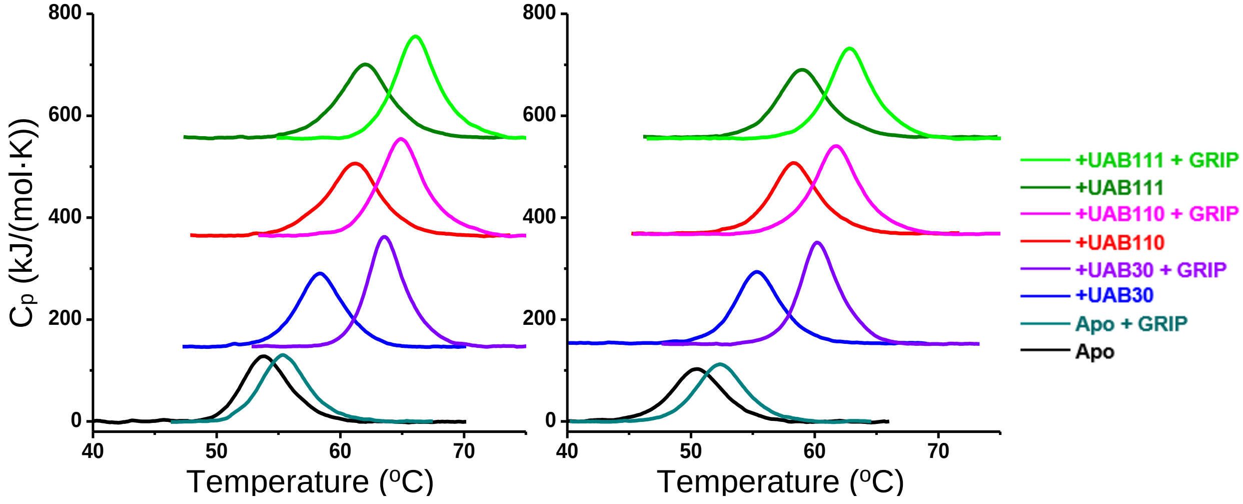


DSC molar heat capacity profiles of 1.5 μM RXRα LBD at pH 8.8 (left panel) or pH 9.5 (right panel), in the presence of no rexinoid or GRIP-1 (*Apo*), 0.4 mM GRIP-1 (*Apo* + GRIP), 30 μM UAB30 (UAB30), 30 μM UAB30 and 0.4 mM GRIP-1 (UAB30 + GRIP), 10 μM UAB110 (UAB110), 10 μM UAB110 and 0.4 mM GRIP-1 (UAB110 + GRIP), 10 μM UAB111 (UAB111), or 10 μM UAB111 and 0.4 mM GRIP-1 (UAB111 + GRIP). The scan rate, *v*, was 4.0 ºC/min. All samples contained 1% DMSO.

**Figure S9. ITC of *apo*-RXRα LBD and UAB30 at different temperatures**

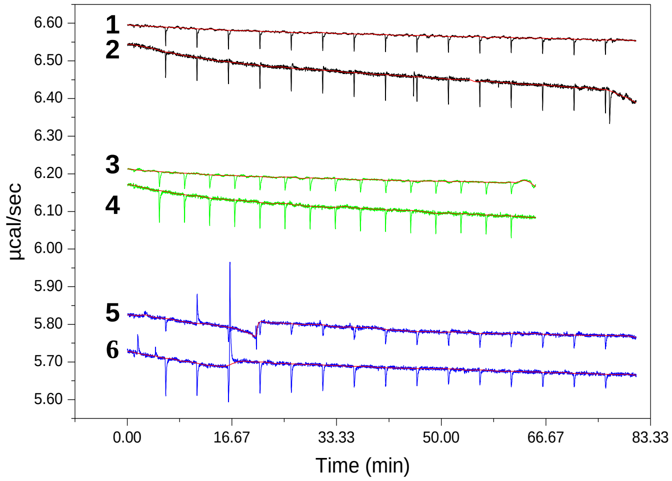


ITC titrations of 20 μM *apo*-RXRα LBD homodimer to 5 μM UAB30 at 10 ºC (trace 2), 20 ºC (trace 4), and 30 ºC (trace 6). Traces 1, 3, and 5 represent the mixing heat at each temperature where 20 μM *apo*-RXRα LBD homodimer was titrated to the same buffer without UAB30.

**Figure S10. DSF $T_m$ of UAB30:RXRα LBD as a function of GRIP-1 concentration**

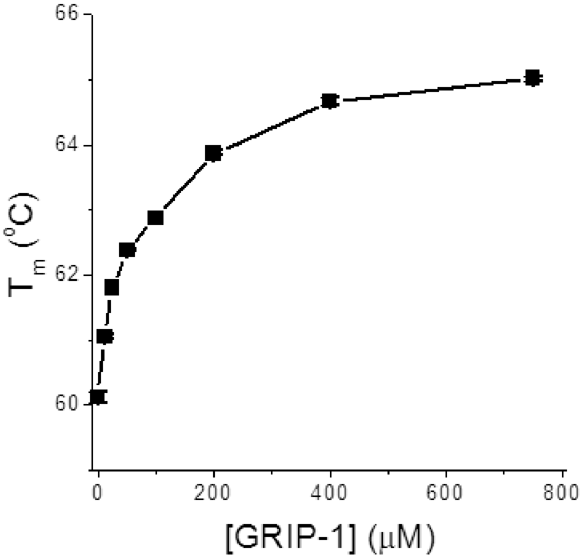


DSF $T_m$ of 2.5μM RXRα LBD homodimer in the presence of 15 μM UAB30 and increasing GRIP-1, determined from the first derivatives of fluorescence at 350 nm as a function of temperature.

**Table S3. Summary of ITC measurements of GRIP-1 to RXRα LBD:rexinoid complexes**

| Temperature (°C) | $K_d$ (μM) | $\Delta H_a$ (kJ/mol) | $-T\Delta S_a$ (kJ/mol) | $\Delta G_a$ (kJ/mol) | $n$ | $\Delta C_p{}^a$ kJ/(mol·K) |
|---|---|---|---|---|---|---|
| **UAB110** | | | | | | |
| 10 | 3.8 | -11.7 | -18.4 | -29.3 | 1.05 | |
| 20 | 3.3 | -23.4 | -7.5 | -30.9 | 1.10 | |
| 30 | 5.6 | -35.1 | 4.6 | -30.5 | 1.18 | -1.19 ± 0.01 |
| **UAB111** | | | | | | |
| 15 | 7.9 | -11.7 | -16.3 | -28.0 | 1.16 | |
| 20 | 2.4 | -18.4 | -13.4 | -31.8 | 1.05 | |
| 30 | 3.2 | -27.6 | -4.2 | -31.8 | 1.25 | -1.11 ± 0.07 |
